# A Gulf Stream frontal eddy harbors a distinct microbiome compared to adjacent waters

**DOI:** 10.1101/2023.02.23.529726

**Authors:** Jessica L. Gronniger, Patrick C. Gray, Alexandria K. Niebergall, Zackary I. Johnson, Dana E. Hunt

**Affiliations:** Marine Laboratory, Duke University, Beaufort NC USA; Nicholas School of the Environment, Duke University, Durham NC USA; Biology and Civil & Environmental Engineering, Duke University, Durham NC USA

**Author notes:** To whom correspondence should be directed: Dana Hunt 135 Duke Marine Lab Rd Beaufort NC 28516.

## Abstract

Mesoscale oceanographic features, including eddies, have the potential to alter productivity and other biogeochemical rates in the ocean. Here, we examine the microbiome of a cyclonic, Gulf Stream frontal eddy, with a distinct origin and environmental parameters compared to surrounding waters, in order to better understand the processes dominating microbial community assembly in the dynamic coastal ocean. Our microbiome-based approach identified the eddy as distinct from the surround Gulf Stream waters. The eddy-associated microbial community occupied a larger area than identified by temperature and salinity alone, increasing the predicted extent of eddy-associated biogeochemical processes. While the eddy formed on the continental shelf, after two weeks both environmental parameters and microbiome composition of the eddy were most similar to the Gulf Stream, suggesting the effect of environmental filtering on community assembly or physical mixing with adjacent Gulf Stream waters. In spite of the potential for eddy-driven upwelling to introduce nutrients and stimulate primary production, eddy surface waters exhibit lower chlorophyll *a* along with a distinct and less even microbial community, compared to the Gulf Stream. At the population level, the eddy microbiome exhibited differences among the cyanobacteria (e.g. lower *Trichodesmium* and higher *Prochlorococcus*) and in the heterotrophic alpha Proteobacteria (e.g. lower relative abundances of specific SAR11 clades) versus the Gulf Stream. However, better delineation of the relative roles of processes driving eddy community assembly will likely require following the eddy and surrounding waters since inception; additionally, sampling throughout the water column could better clarify the contribution of these mesoscale features to primary production and carbon export in the oceans.

## Introduction

Marine microorganisms are of the engines of biogeochemical cycles, supporting critical ecosystem functions on a global scale; yet these biogeochemical processes depend on the composition, function, and activity of microbial communities (1). Marine microbiology has emphasized environmental filtering (how environmental factors determine microbiome composition due to bottom up effects), in determining community assembly, including factors such as water temperature, salinity and nutrient availability (2, 3). However, processes affecting microbiome community assembly include biological interactions, dispersal limitation and stochasticity (4), as well as the microbiome’s ecological history (5). These historical contingencies, the effects of prior conditions on current microbiome assemblages, have emerged as a significant force shaping communities (6, 7). For example, prior environment changes can result in microbiomes that are more resistant (composition remains unchanged after the disturbance) or resilient (composition is altered but eventually returns to pre-disturbance state) to future perturbations (8, 9). Similarly, microbiomes from stable environments may be more sensitive to environmental changes (9, 10). However, in aquatic ecosystems, establishing antecedent environmental conditions is particularly challenging due factors including water movement, unmeasured disturbances and environmental parameter seasonality (11–13). Further, although dispersal limitation is assumed be reduced in aquatic systems compared to other systems, such as soils, oceanographic features such as steep frontal gradients between water parcels or vertical stratification can limit dispersal (14), and thus constrain microbiome composition. Here, we examine a frontal eddy, which may be able to isolate and transport microbial populations (15), to investigate the relative roles of environmental filtering, ocean physics and historical contingencies in an aquatic system.

Frontal eddies form along western boundary currents when instabilities in the current, often stemming from interactions with bathymetry, cause meanders to form and evolve into distinct cyclonically rotating water parcels. While considered mesoscale eddies, frontal eddies are smaller and less stable than typical mesoscale eddies such as Gulf Stream rings (16). Gulf Stream rings form downstream from Cape Hatteras once the current is freely meandering and can “pinch off”, in contrast, frontal eddies do not detach from the Gulf Stream and stay trapped in the meander of the current. The eddy core is shelf water that is entrained in this Gulf Stream meander and a shallow warm streamer flows from the downstream meander crest around this core, separating the eddy from the shelf (17). All frontal eddies are cyclonic, and thus rotation typically uplifts colder, nutrient-rich waters into the euphotic zone, often stimulating phytoplankton growth, though continued uplift is dependent on the eddy’s age (16–18). Beyond frontal eddies, there has been substantial oceanographic interest in eddies generally, particularly, their roles in temporal and spatial variability in primary production, carbon export to the deep sea, and rapid shifts in marine microbiomes (19–21). Ecologically, eddy formation partially isolates waters, with the potential to develop microbiomes, environmental parameters, and biogeochemical rates distinct from surrounding environments (22, 23). However, it is difficult to predict eddy microbiomes; the specific microbial taxa stimulated vary among the phytoplankton and include diatoms, prasinophytes, dinoflagellates, and *Synechoccocus* (24–26), with the potential for concurrent increases in multiple phytoplankton (26), and with likely consequences for heterotrophic composition and organic matter export (24, 25, 27). Differences among eddy microbiomes have been attributed to eddy characteristics, such as the initial microbiome, environmental factors (temperature, nutrient fluxes, mixed layer depth, etc.), and the eddy stage, among others (24, 28). Thus, although mesoscale eddies are pervasive throughout the global ocean, and frontal eddies are ubiquitous along western boundary currents, we have poor predictive ability of their influence on microbiomes and associated biogeochemical processes.

This study takes place just northeast of Cape Hatteras, NC, USA (Figure 1A) which is a dynamic point where the warm, salty, rapidly-flowing Gulf Stream, the cooler and fresher Slope Sea, and the even colder and fresher mid-Atlantic Bight shelf water meet, and where the Gulf Stream separates from the continental shelf. The study area is a region of high gradients, moving from the productive continental shelf waters to the oligotrophic open ocean in a few dozen kilometers across shore. Here, we compare a Gulf Stream frontal eddy’s microbiome with those of the adjacent continental shelf, continental slope, and Gulf Stream waters off the coast of Cape Hatteras. Gulf Stream frontal eddies form ∼every 3-7 days in the South Atlantic Bight (18), can occupy nearly half of the front between the Gulf Stream and the continental shelf in this region (29), and have been shown to play a significant role in nutrient transport into the euphotic zone, enhancing primary and possibly secondary production both in the eddy and on the continental shelf (17, 23, 30). Using a combination of field and satellite-based (temperature and chlorophyll *a*) measurements, we take a microbiome-centric approach to identify water parcels and population-level differences between water parcels in order to better understand the forces shaping the microbiomes of these mesoscale ocean features.

**Figure 1.**
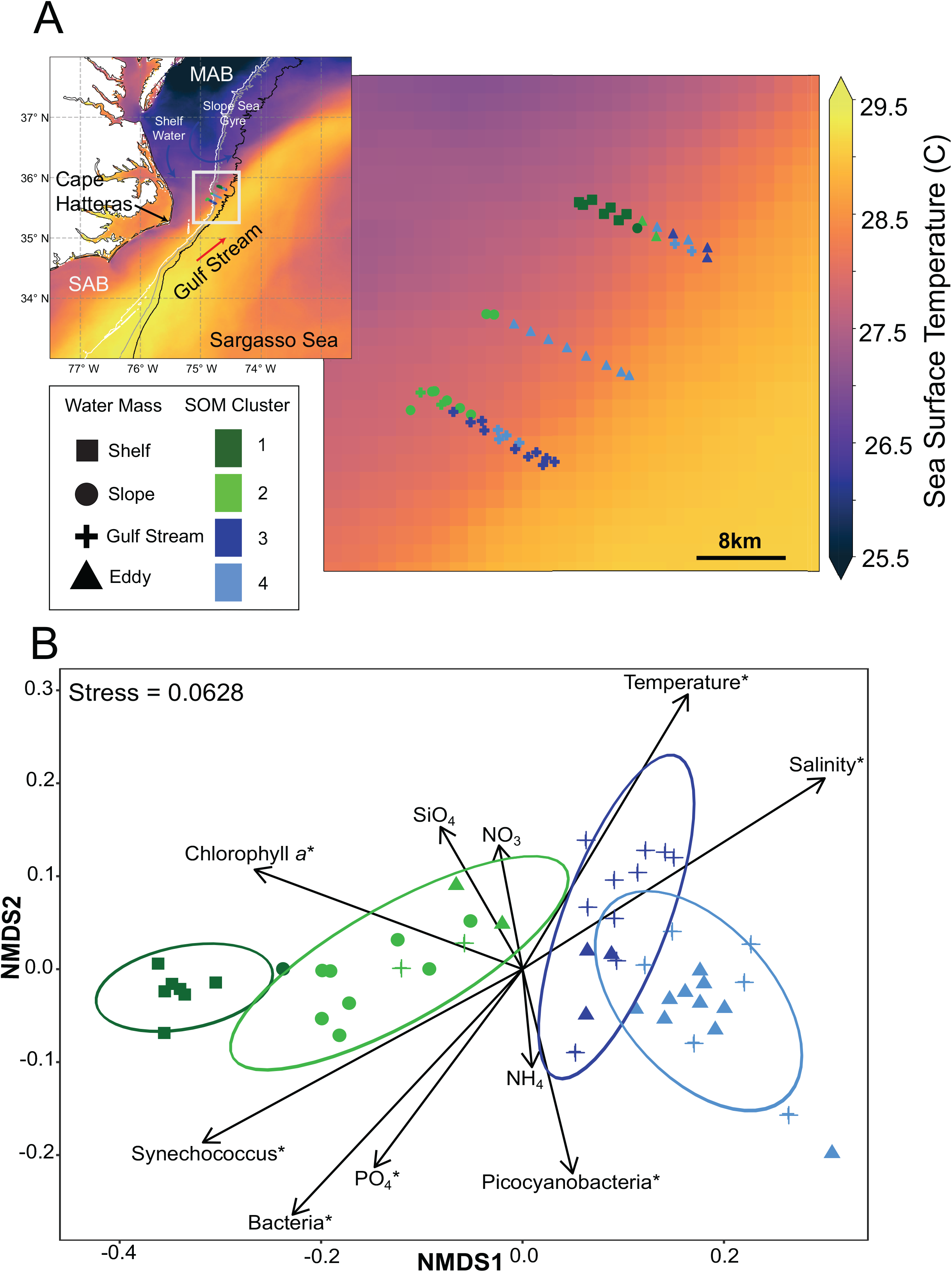
Locations and microbiome of field samples. (A) Map of sampling sites for five ∼10- 15 km long ship-board transects off the coast of Cape Hatteras, North Carolina, USA September 5-7, 2021. Background color represents average sea surface temperatures from August 20^th^ to September 10^th^ obtained from the Geostationary Operational Environmental Satellite 16 (GOES- 16). Each sampling point is shown along the transect and categorized by temperature and salinity (shapes) and microbial-community based k-means clusters (colors), generated using the self-organizing map (SOM) R package ‘kohonen’. Isobaths for 500m (white), 1000m (grey) and 2000m (black) are included for context. (B) Non-metric multidimensional scaling (NMDS) ordination computed based on Bray-Curtis dissimilarity for 16S rRNA gene libraries. Ellipses show the multivariate t-distribution 95% confidence interval for the mean of each SOM cluster. Environmental factors found to be statistically significant (Multiple regression permutation test, p<0.05) are indicated with an ‘*’.

## Results & Discussion

Using satellite data, we followed a single Gulf Stream frontal eddy from formation on the Charleston Bump until sampling ∼2 weeks later (Figure 2). This sampled Gulf Stream frontal eddy was visible via satellite as a cooler, chlorophyll *a* enriched region relative to the surrounding Gulf Stream waters (31); however, there was no evidence of a surface bloom or enhancement of chlorophyll *a* levels compared to the shelf waters. In fact, satellite- measured surface chlorophyll *a* levels decline over time, likely either through nutrient depletion or mixing with Gulf Stream waters (Figure 2)(31). Here, we *in situ* sampled the eddy (and surrounding waters) as it moved past Cape Hatteras on September 5-7^th^ 2021. At this point the eddy was approximately 20 by 50 km (longer in the along stream dimension) and we collected five across-stream transects of 10-15 km each (49 stations with 1-2 km between stations) to obtain high-resolution measurements of surface microbial community composition and environmental conditions (Figure 1A).

**Figure 2.**
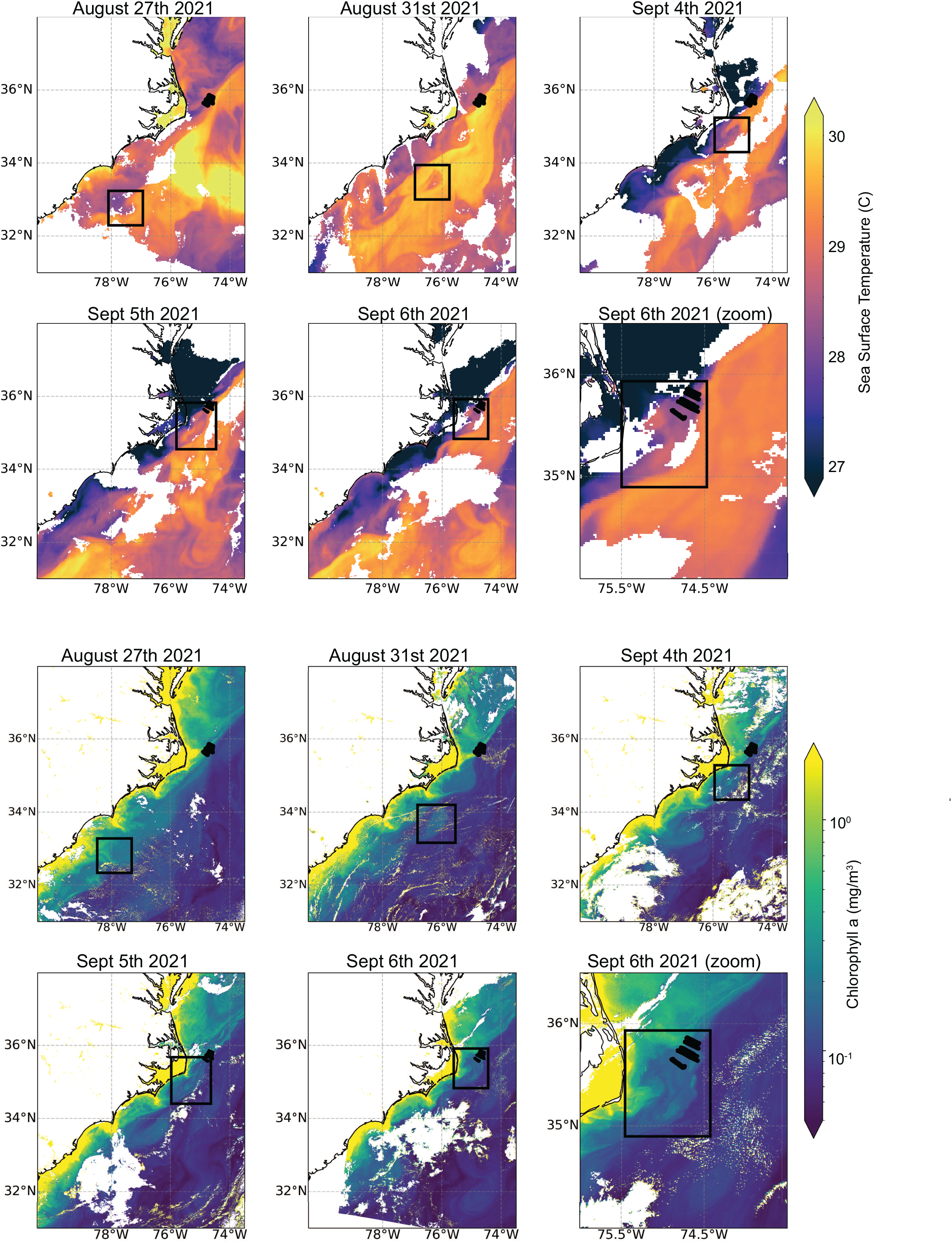
**Satellite overview of the observed frontal eddy from formation to *in situ* sampling**. Black dotted lines are all 5 transects from this study. Black boxes outline the eddy location across five days from August 27th to September 6th. SST is from GOES-16 and chlorophyll *a* is from Sentinel-3’s OLCI sensor. Boxes were determined manually by identifying an anomaly in SST and ocean color measurements that is consistent with a cyclonic frontal eddy and associated warm streamer.

Initially, surface temperature and salinity were used to identify water parcels (Figure S1) (31). However, these physical parameter-based categories did not align with microbiome composition, particularly among the Gulf Stream and eddy samples (Figure S2A). Thus, we used a microbiome-based, self-organizing map (SOM) approach (32) to cluster the microbiome data independently of environmental variables. We then compared these biologically-based clusters (SOM) with the temperature and salinity-based water parcel classifications. Although these two schemes did overlap, there were also a number of disagreements in the assignments (Figure 1B): SOM cluster 1 largely corresponds to the continental shelf (87.5%), cluster 2 the continental slope (67%), cluster 3 the Gulf Stream (77%) and cluster 4 the eddy (62.5%) stations. Satellite data suggests that disagreements between these simple physical parameters and microbiome-based classifications occurred at transitions between water parcels, particularly around the periphery of the eddy (Figure S3), which could be due to complex mixing between the eddy and surrounding Gulf Stream. We found that the microbiome-based (SOM) approach classified more samples as eddy-associated. Therefore, identification of the eddy based on temperature and salinity alone could underestimate the geographic extent of the eddy’s biogeochemical influence (Figure S3), or fail to identify mixing of the eddy microbiome into surrounding waters.

Henceforth, for clarity, these statistically-distinct SOM clusters (ANOSIM, p<0.05) will be referred to by their most common location (1 = continental shelf, 2 = continental slope, 3 = Gulf Stream, 4 = eddy), with the assumption that SOM clusters most accurately map potential ecological differences between the samples. However, disagreements between clustering based on surface properties (temperature, salinity) and microbiomes merits further investigation in both the biological and physical oceanographic fields, particularly at eddy margins to better identify microbiome origins and implications (e.g. changes in biogeochemical rates).

Next, we examined how environmental factors differed among these clusters; the shelf and slope clusters are also clearly distinct on the NMDS, with relatively higher overlap for the Gulf Stream and eddy clusters (Figure 1B). We examined environmental variables that might help explain the observed microbiome patterns, the continental shelf and slope exhibit higher cell abundances (total prokaryotes, *Synechococcus*) and higher PO_4_ along with lower salinity and cooler water temperatures compared to the eddy and Gulf Stream (Figures 3, S4; Pairwise Wilcoxon ranked sum test p < 0.05). This is generally consistent with higher resource availability on the continental shelf and slope compared to the Gulf Stream and eddy, indicating greater terrestrial influence and resources in near-shore environments and oligotrophic conditions beyond the shelf break (33). Ordination of the microbiome in relation to select environmental variables [temperature, salinity, chlorophyll *a*, nutrients (SiO_4_, NO_3_, NH_4_, PO_4_), and flow-cytometric abundances of bacteria and cyanobacteria] identified significant associations for variables other than SiO_4_, NO_3_ and NH_4_, (Figure 1B, Table S1), highlighting the factors which could explain the microbiome composition. Here, we specifically focused on comparison of the Gulf Stream and eddy samples as they are most similar in both environment and microbiome: the eddy is ∼1 °C cooler, has lower chlorophyll *a*, lower *Synechococcus* and higher non-phycoerythrin containing picocyanobacteria (hereafter labeled *Prochlorococcus*) concentrations (Figures 3, S4; Pairwise Wilcoxon ranked sum test p< 0.05). While PO_4_ in the eddy is statistically higher than in the Gulf Stream, eddy phosphate concentrations have a large range, suggesting potential variability in nutrient fluxes/ availability within this water parcel (Figure 3D). Thus, in contrast to other cyclonic eddies (23, 34), here, eddy surface waters generally exhibit oligotrophic characteristics (low nutrients, high *Prochlorococcus* concentrations, lower chlorophyll *a,* but higher PO_4_) compared to the Gulf Stream, in spite of forming via entrainment of presumably higher nutrient continental shelf water and the potential for upwelling-driven nutrient additions (Figures 3, S4). Overall, these results are consistent with the satellite measurements that do not show enhanced surface photosynthetic biomass of the eddy. Possible explanations for this result include that photosynthetic biomass enhancement occurred primarily below the mixed layer and was not visible at the surface, or that high stratification in the summer along with a deep euphotic depth prevented the eddy-associated uplift from enhancing productivity. Examination of microbial community characteristics (diversity etc.) as well as specific phylotypes associated with each water parcel may offer additional clues to factors that shaped these communities in the coastal ocean.

**Figure 3.**
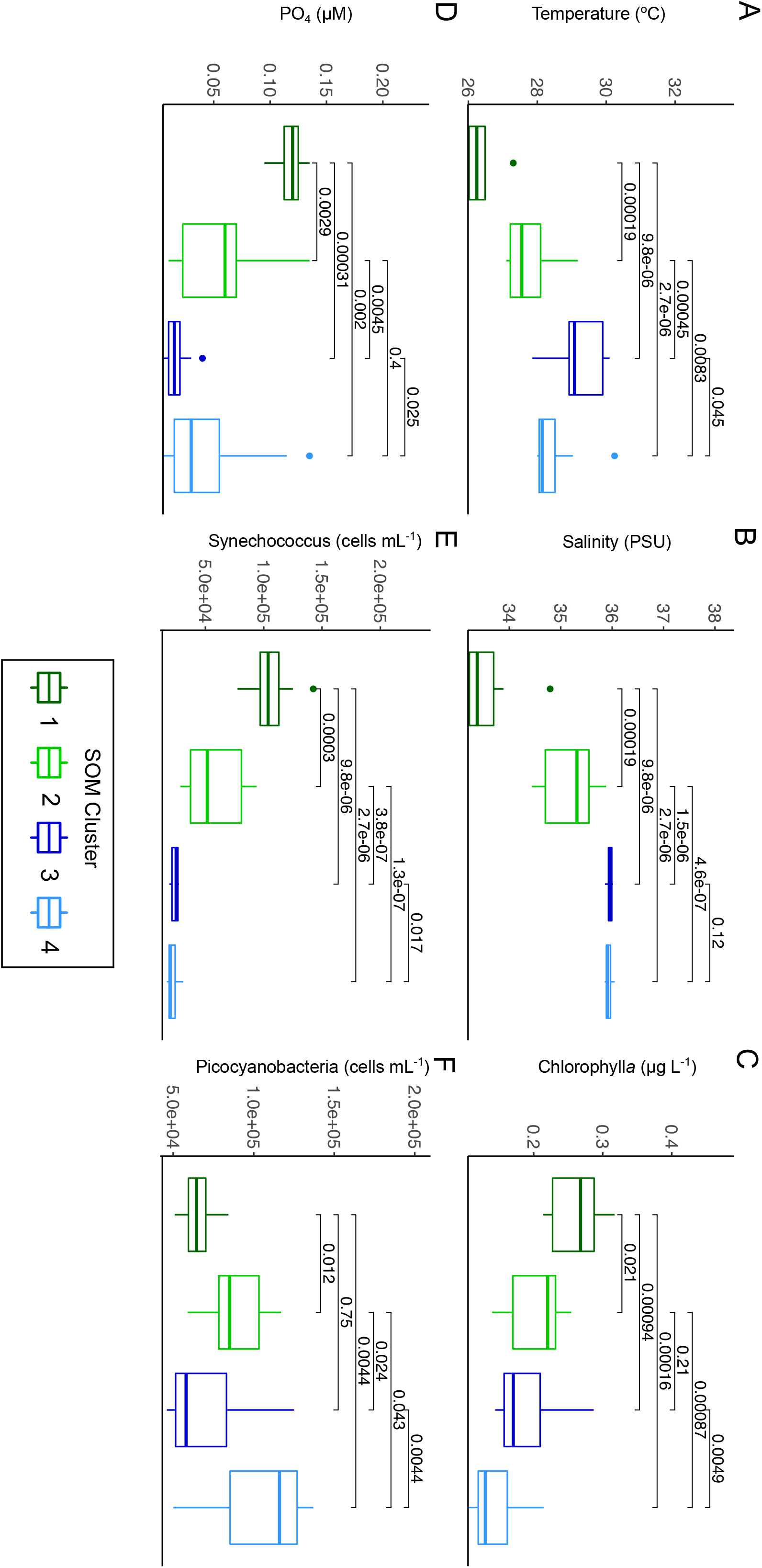
Key environmental factors for Self-Organizing Map (SOM) clusters Numbered clusters correspond to the most common environments: Continental shelf (Cluster 1), Continental slope (Cluster 2), Gulf Stream (Cluster 3), Eddy (Cluster 4). For the box and whisker plots, the center lines denote the median value while the box encompasses the 25^th^ to 75^th^ percentiles of the dataset. Whiskers denote the 5^th^ and 95^th^ percentiles and values that fall outside these ranges are shown as individual points. Brackets indicate p-values of Wilcoxon Rank Sum pairwise comparisons. Additional environmental parameter comparisons are shown in Figure S4.

### Microbial Assemblages

As eddies can alter microbial community parameters, potentially increasing diversity through mixing of water parcels and influx of resources, or conversely reducing diversity and evenness through competitive exclusion (20, 35–37), we compared diversity and evenness indexes in the four water parcels. Notably, the eddy exhibits lower richness and evenness compared to the Gulf Stream (Figure 4; Wilcoxon ranked sum test p<0.05), reflecting the dominance of specific taxa, particularly *Prochlorococcus* (35) (Figure 5). While the eddy initially formed from continental shelf waters (although off the coast of South Carolina) and retains comparable richness to shelf waters in these transects (Figure 4), the eddy is statistically less even than the other water parcels ∼two weeks after formation, suggesting continued development of the microbiome (Wilcoxon ranked sum test p<0.05). While the eddy microbiome is most similar to the Gulf Stream’s, if dispersal limitation significantly shapes the eddy microbiome and if we assume the shelf community where the eddy formed is similar to the shelf community off Cape Hatteras, we might predict that presence-absence data (Sorenson index) would group the eddy with the continental shelf and that relative abundances would shift to match the current environmental conditions. However, the Sorenson index also groups the eddy and the Gulf Stream samples (Figure S5), as was observed for Bray-Curtis dissimilarity (Figure 1B), suggesting that environmental filtering or physical mixing shapes frontal eddy communities.

**Figure 4.**
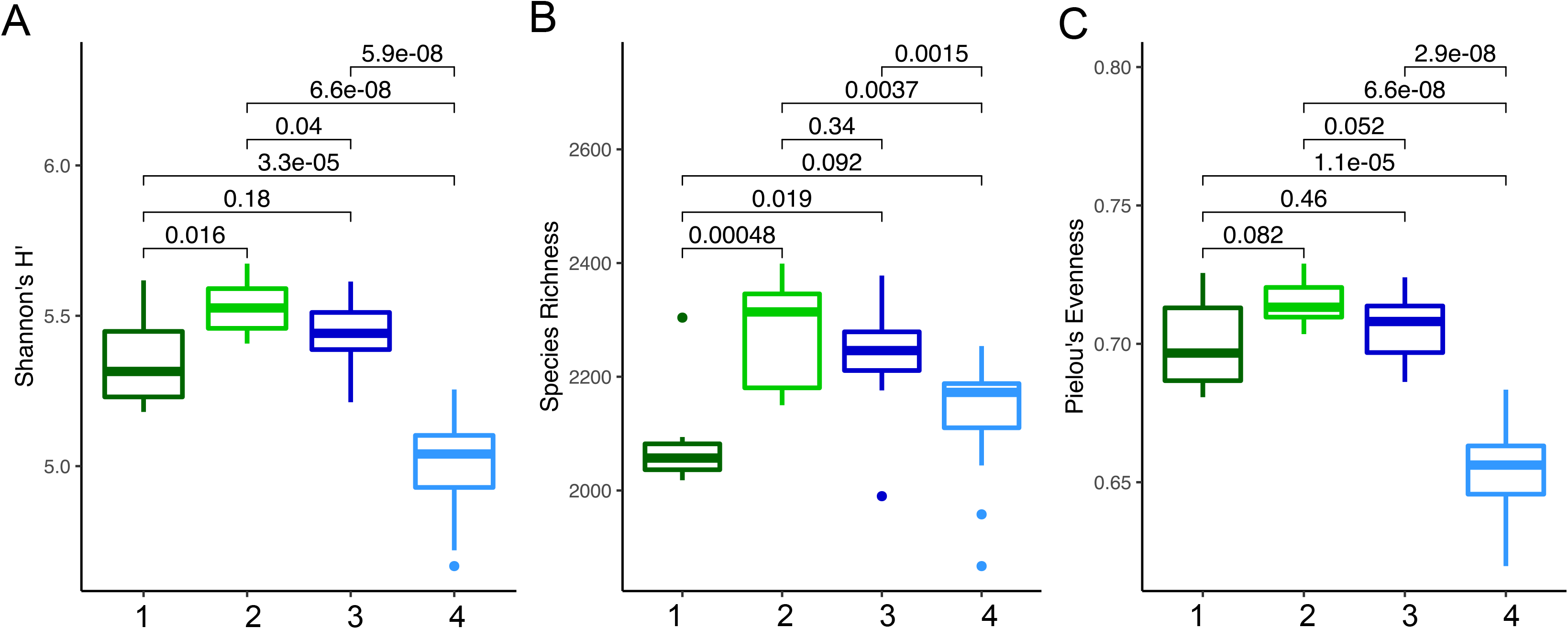
**Alpha diversity metrics for 16S rRNA gene community composition for each Self-Organizing Map (SOM) cluster**. (A) Shannon’s H index, (B) species richness, and (C) Pielou’s evenness were calculated based on the absolute abundance of ASVs using ‘vegan’ (v2.6.2) in R (v4.0.0). SOM cluster numbers correspond to the following classifications: 1 –continental shelf, 2-continental slope, 3- Gulf Stream, 4- eddy. Brackets indicate p-values associated with Wilcoxon Rank Sum test pairwise comparisons.

**Figure 5.**
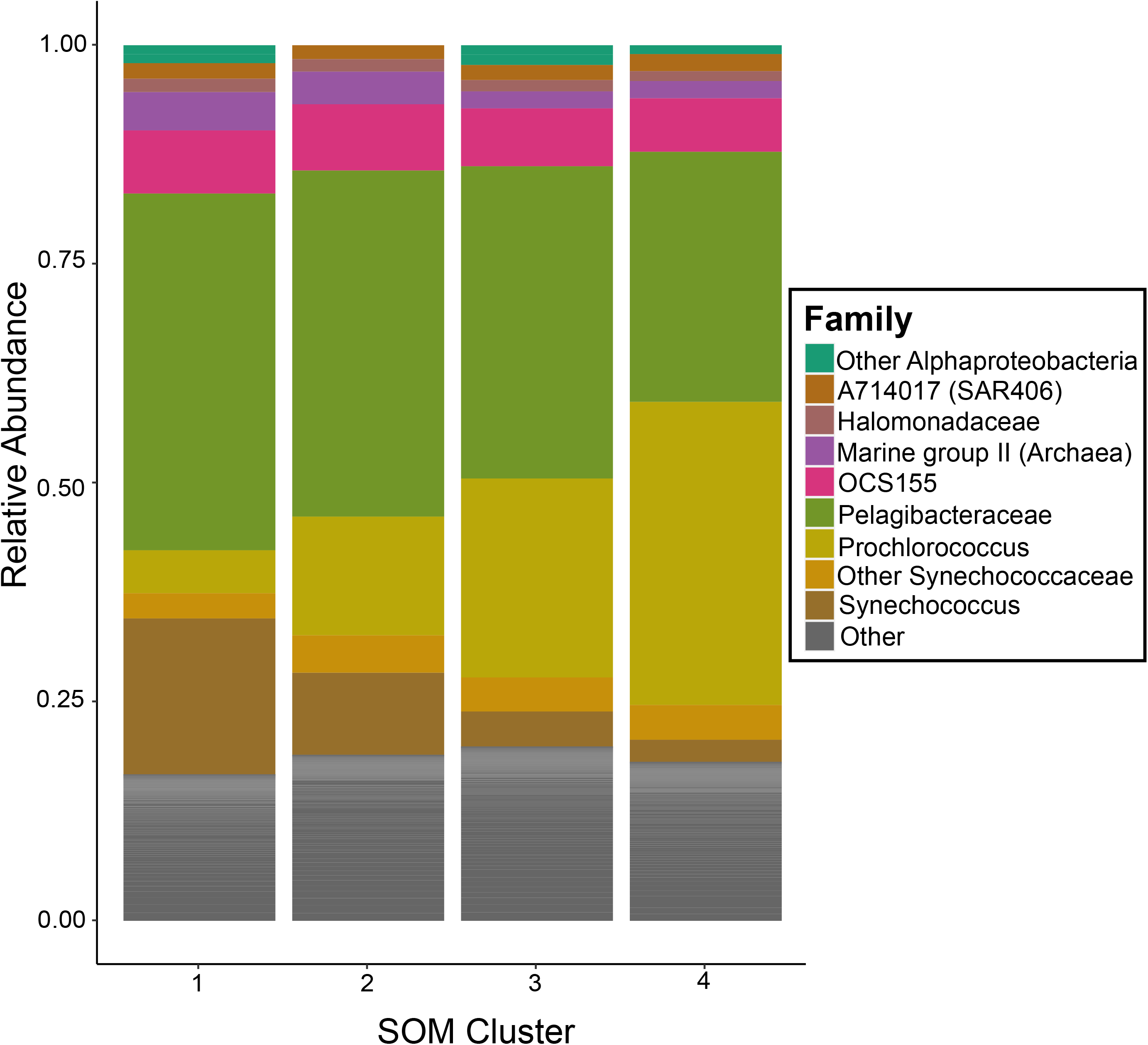
**Average family-level composition for each Self-Organizing Map (SOM) cluster**. The average community was calculated by taking the mean relative abundance of each taxon across all samples within that cluster, then normalizing relative abundances to 100%. ASVs with a relative abundance of less than 1% are grouped as “Other.” Amplicon Sequence Variants (ASVs were generally grouped at the family level, but when family is undefined they are labeled at the phylum level. Additionally, the family *Synechococcaceae* was resolved at the genus level to illustrate the *Synechococcus*-*Prochlorococcus* inversion between clusters.

Across all samples, the microbiome composition reflects abundant coastal microbes including *Synechococcaceae* (∼25-40%), *Pelagibacteraceae* (∼28-40%), Actinobacteria family OCS155 (∼7%), the Archaeal Marine Group II clade (∼2-4%) and the SAR406 clade A714017 (∼2%; Figure 5) (3, 33, 38, 39). Yet, we observed onshore-offshore spatial transitions (33), including the switch within *Synechococcaceae* genera, with *Synechococcus* dominating the shelf and slope, and *Prochlorococcus* the Gulf Stream and eddy (Figure 5). As specific taxa may offer additional clues about the factors which shaped these microbiomes, we examined ASV relationships with specific clusters, using 250 most abundant ASVs (representing ∼73% of the microbial community) with both linear discriminant effect size analysis (LEfSe) and a Bayesian generalized joint attribute model (GJAM) (40, 41) (Tables S2, S3). Both LEfSe and GJAM identify taxa associated with specific water parcels (Figure 6); moreover, these approaches generally agree: identifying increased relative abundance of *Pelagibacter* and *Synechococcus* in continental shelf and slope waters and enrichment of *Prochlorococcus* and *OCS155* in Gulf Stream and eddy waters, which generally aligns with expectations of oligotrophic, warm water-associated taxa offshore (33). GJAM identified a single difference between the eddy and Gulf Stream associated taxa, a High Light clade I *Prochlorococcus* ASV (Tables S2,S3; Figure 6B) that was enriched in both the eddy and slope samples, potentially suggesting either a continued legacy of nearshore waters or mixing of slope waters into the eddy (Figure 6B). Although there are similarities between the eddy and Gulf Stream samples, their differences help identify the current (e.g. environmental filtering) and antecedent (e.g. historical contingencies) conditions that shape the microbiome.

**Figure 6.**
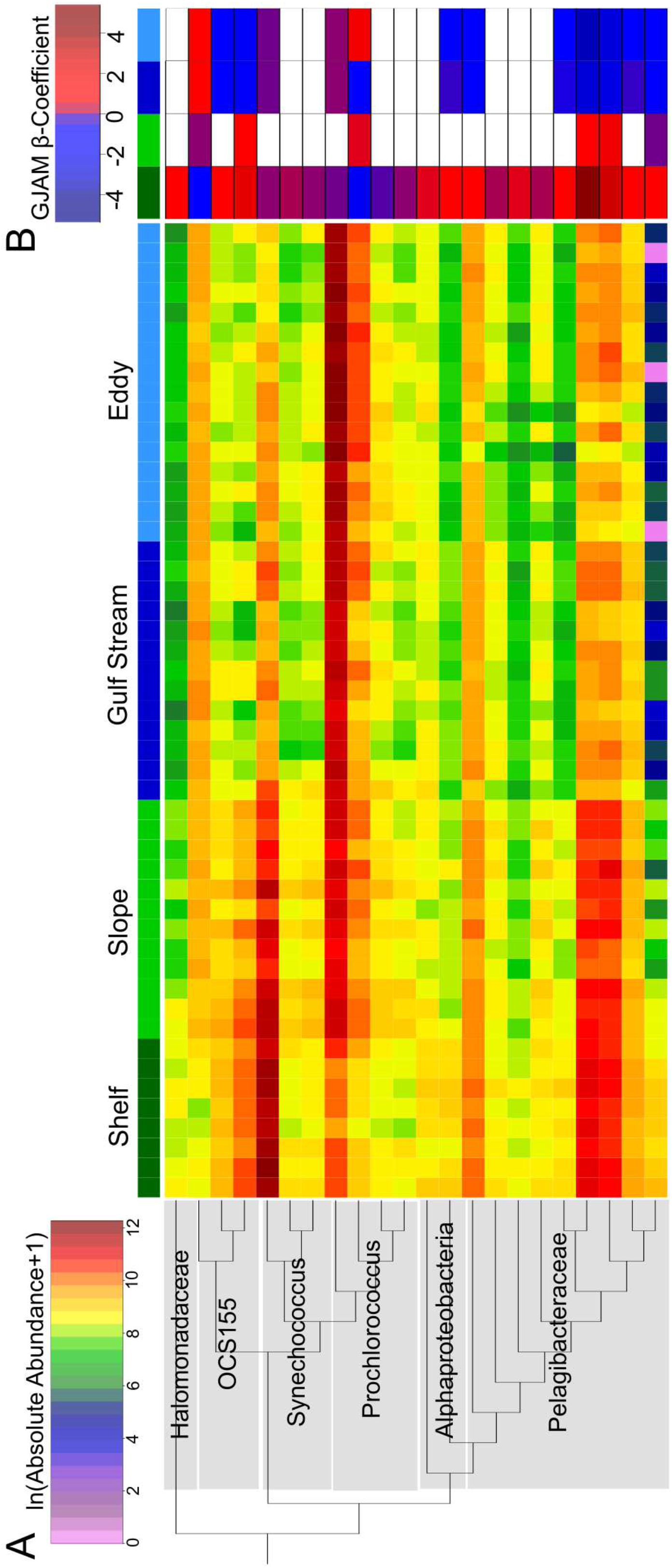
Amplicon sequence variant (ASV) associations with specific SOM groupings. (A) Heatmap of ln(absolute abundance+1) of the 22 ASVs identified with at least one significant SOM cluster beta coefficient obtained from generalized joint attribute modeling (GJAM) analysis of the 250 most abundant ASVs across all samples. Samples are organized by SOM grouping (labeled at top) (B) Heatmap of statistically significant SOM cluster-associated beta coefficients (95% confidence interval does not overlap 0), non-significant beta coefficients are shown as white. Colored labels at the top of each column identifies the SOM cluster (as in panel A). ASVs are organized maximum likelihood phylogenetic tree constructed using Smart Model Selection in PhyML 3.0 with taxonomic assignments based on the RDP naïve Bayesian classifier using the Greengenes version 13.5 database.

A comparison of just the eddy and Gulf Stream using both LEfSe and DESeq2 (42) more clearly reveals distinctions between their microbiomes (Table S4; Figure 7). A number of SAR11 lineages were enriched in the Gulf Stream compared to the eddy (Figure 7). The SAR11 clade is known to contain streamlined heterotrophs (43); however, we can only speculate that ASV differences could be due to resource differences or stochastic effects. Several high-light clade *Prochlorococcus* ASVs were enriched in the eddy over the Gulf Stream which is consistent with flow-cytometric non-phycoerythrin containing picocyanobacteria enrichment in the eddy (Figure 3F) and their role as efficient nutrient recyclers (44). Interestingly, although inorganic nutrients are low in both the eddy and Gulf Stream (Figures 3, S4), the Gulf Stream samples are enriched in a putatively nitrogen-fixing cyanobacterium *Trichodesmium* ASV (Figure 7) (45). Thus, within the cyanobacteria and despite similar inorganic nitrogen concentrations, the Gulf Stream and eddy microbiomes may reveal differences in nutrient fluxes or the role of historical contingencies in microbiome composition (46). Moreover, we acknowledge that standing stocks (e.g. nutrient concentrations) are not equivalent to nutrient turnover rates, as resources can be rapidly cycled in oligotrophic conditions (47). The development of the colonial, nitrogen-fixing *Trichodesmium* may require additional time or lower nitrogen fluxes. As the eddy sustains higher concentrations of *Prochlorococcus* and total bacterioplankton, and statistically higher levels of PO_4_, this data could indicate that the eddy has experienced higher nutrient fluxes, which could support higher primary and secondary production, although this is not evident in standing photosynthetic biomass (as measured by chlorophyll *a*, Figure 3C). Although there was no satellite-detectable eddy surface bloom, and the eddy microbiome likely shifted toward that of the Gulf Stream over time, microbial communities can offer clues about the processes occurring in these dynamic mesoscale features.

**Figure 7.**
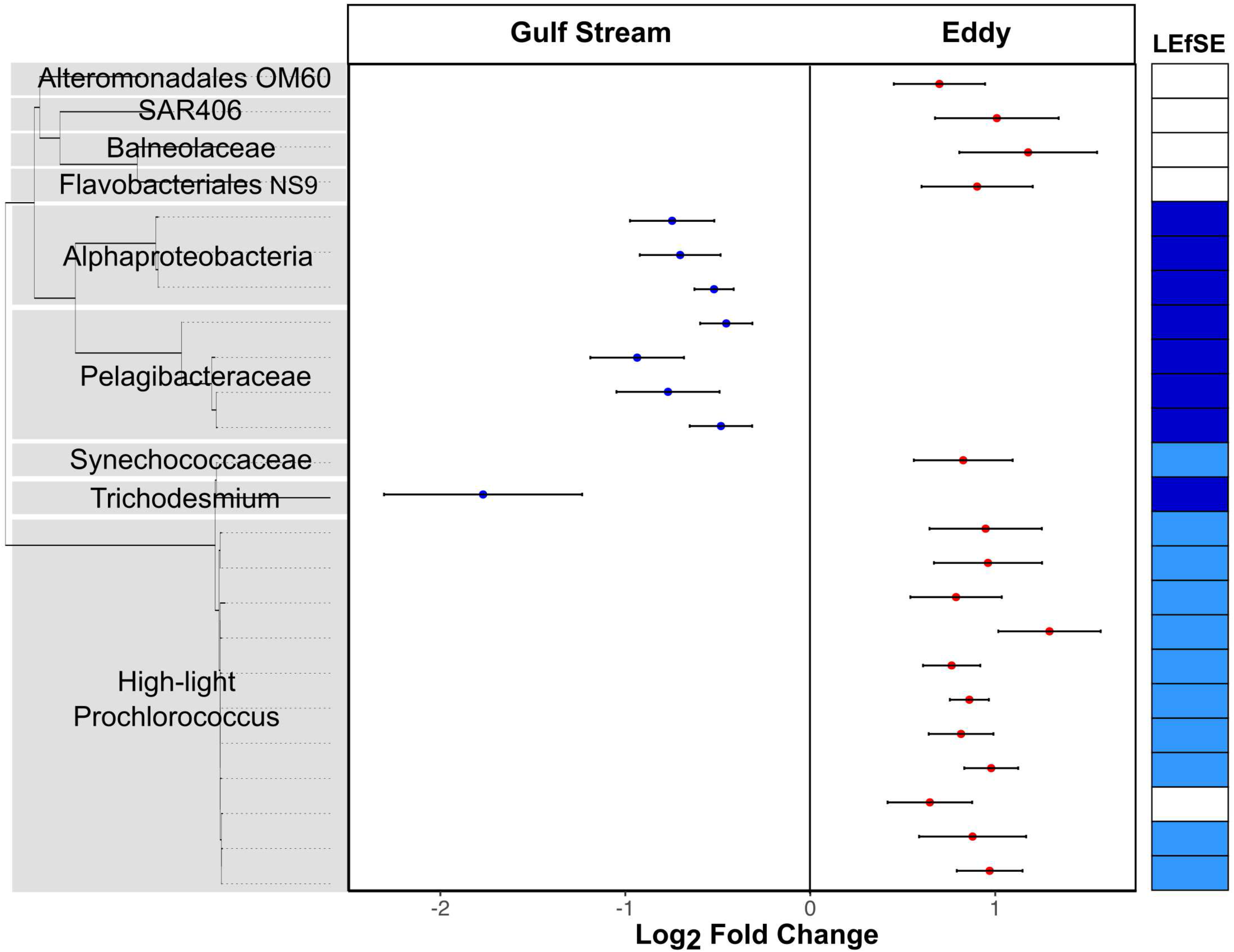
Microbiome differences between the Gulf Stream and eddy. Left panel: Phylotypes (ASVs) that exhibited significant differential abundance in the eddy relative to the Gulf Stream as identified using DESeq2, left side indicates enrichment in the Gulf Stream, right side eddy.

Dots indicates the Log_2_ fold changes for each ASV identified as significant (Benjamin-Hochberg multiple hypothesis adjusted p<0.05), error bars indicate the standard error of the Log_2_ fold changes as calculated by DESeq2. Only ASVs with a minimum average relative abundance of 0.05% across the samples being compared were included in the analysis. Y-axis represents maximum likelihood phylogenetic tree constructed using Smart Model Selection in PhyML 3.0. Taxonomic assignments were obtained from the RDP database. Right hand column indicates those ASVs that were also found to have a significant association with either the Gulf Stream (dark blue) or the eddy (light blue) as identified using Linear discriminant analysis Effect Size (LEfSE).

## Conclusions

Here we show that a cyclonic, frontal eddy that formed along the western boundary of the Gulf Stream harbored a microbial community distinct from those of surrounding waters. This eddy community appears to be shaped by either environmental selection or mixing with the Gulf Stream (16, 29), rather than dispersal limitation, as it was most similar to that of the Gulf Stream rather than continental slope waters where it originated. While this study was limited to surface samples and encompasses a brief period in the eddy’s development, new technologies are enabling Lagrangian sampling of eddy parameters during development and throughout the water column to better quantify upwelling, eddy coherence and mixing with the Gulf Stream, across depths (48–51). High-resolution, *in situ* sampling will enable the field to develop a more quantitative assessment of how source populations and environmental drivers interact to shape these dynamic mesoscale features over time. Although more research is needed to understand the impact of eddies globally, this study deepens our understanding of the forces shaping Gulf Stream frontal eddy microbiomes off the South Atlantic Bight and provides important directions for future studies. While there was not support for this eddy contributing to the biodiversity hotspot located off of Cape Hatteras (36, 52), other frontal eddies may enhance biodiversity by delivering distinct microbial communities, enhanced zooplankton populations, and upwelling of resources (31). Furthermore, the frequency of these eddies and their predicted nutrient injections into the Gulf Stream may play a significant role in carbon sequestration and productivity along the South Atlantic Bight. Although we found this eddy harbored a distinct microbial community, further investigations will be needed to better understand the role of historical contingencies and the temporal dynamics of eddy-associated communities and their potential role in global biogeochemical cycles. Moreover, this approach of microbially-defined, environmental variable agnostic sample clustering can be applied more broadly in oceanography to help disentangle potentially ecologically-distinct microbiomes.

## Methods

### Sample collection and sea water parameter characterization

We tracked via satellite a cyclonic, frontal eddy that formed from shelf water off of Charleston, South Carolina, USA on August 25-26^th^ 2021, was isolated by warmer, Gulf Stream-derived waters, and then traveled north along the Gulf Stream’s western front (31). Seawater was collected on board the *R/V Shearwater* on 5 transects from September 5-7^th^, 2022; transects were planned to cross the front between the shelf and eddy or shelf and Gulf Stream waters. Water samples were collected during transit from the ship’s flow-through seawater system (∼1 m depth). In general, the transects were 10-15 km long (over 2-3 hours) with 14-21 samples collected per transect at approximately 10 min intervals for a total of 86 sampling points, 49 of which included filtration for later nucleic acid analysis. Temperature and salinity were measured with a SeaBird SBE38 Thermosalinograph. Chlorophyll *a* concentrations were measured by filtering 100 mL of seawater onto a ∼0.7 µm glass fiber filter (Whatmann APFF02500) using gentle (<10mm Hg) vacuum. Chlorophyll *a* pigments were extracted in 100% methanol for 48h at -20 °C and fluorescence was measured using a Turner Designs 10-AU fluorometer as previously described (53). Duplicate whole seawater samples for flow-cytometric analyses were collected, fixed with net 0.125% glutaraldehyde and stored at -80 °C until processing.

Prokaryotic phytoplankton populations were enumerated using a Becton Dickinson FACSCalibur Flow Cytometer and categorized as previously described (53). Bacterioplankton were quantified using SYBR Green-I on the Attune NxT acoustic flow cytometer (Life Technologies) (54). Duplicate 0.22 µm Sterivex (Milipore) filtered seawater samples were collected for dissolved inorganic nutrient quantification and stored at -80 °C until processing. Nutrients (NO_2_, NO_3_, PO_4_, SiO_4_) were analyzed at the UC San Diego Scripps Institution of Oceanography ODF chemical laboratory using a Seal Analytical continuous-flow AutoAnalyzer 3 (55, 56). Detection limits for the nutrients analyzed are as follows: NO_2_ = 20nM, NO_3_ = 20nM, PO_4_ =20nM, SiO_4_ = 200nM. No permits were required for environmental sampling within US coastal waters.

### DNA extraction and library preparation

Microbial DNA was collected by filtering ∼4 liters of whole seawater through a 0.22 micron Sterivex filter (Millipore). Filters were stored at -20 °C while on the research vessel and then at - 80 °C upon return to the lab until extraction. Genomic DNA for 16S rRNA gene libraries was extracted using the Gentra Puregene Yeast/Bacteria kit (QIAGEN) supplemented with 60 seconds of bead beating. Extracted DNA was quantified using a Nanodrop ND-100 and 16S rRNA gene libraries were generated using gene primers targeting the V4-V5 region 515F- (5′- GTGYCAGCMGCCGCGGTAA) and 926R (5′-CCGYCAATTYMTTTRAGTTT) (57, 58).

PCR reactions were performed in triplicate with 20 μL reactions containing 20 ng template DNA, 1× Taq Buffer, 0.5 μM of each primer, 200 μM of dNTPs, and 0.4 U of Q5 DNA polymerase (NEB). The thermal cycling conditions were 30 sec at 98 °C, followed by 25 cycles of 10 sec at 98 °C, 30 sec at 50 °C, 30 sec at 72 °C, and a final extension at 72 °C for 2 min.

Triplicate PCR reactions were pooled and gel purified. Libraries were pooled and sequenced at the Duke Center for Genomic and Computational Biology using V2 2 × 250 bp sequencing on the Illumina MiSeq.

### Sequence Processing

Barcodes were removed and sequences were demultiplexed and assigned to samples using Sabre (https://github.com/najoshi/sabre). Sequences were cleaned and clustered using VSEARCH v2.5.1 (59). Low quality sequence ends were trimmed at the Phred quality score (Q) of 30 using a 10 bp running window. Paired-end reads with 10 bp overlap and no mismatches were merged and sequences with expected errors >1 and/or a length < 360 bp were removed. Minimum entropy decomposition analysis (v2.1) was used to resolve amplicon sequences variants (ASVs) (60). Sequencing-error associated noise was reduced by retaining only those ASVs with unique sequences with a minimum abundance of 20 occurrences. Samples contained an average of 39,936 representative reads. Representative ASV sequences were assigned taxonomy using the RDP classifier in MacQIIME v1.9.1. Mitochondrial sequences were removed and libraries were subsampled to 14,187 reads per library. AmpliCopyrighter (v.0.46) was used to correct for differential gene copy numbers across taxa (61). “Absolute abundance” was calculated by multiplying relative abundances of ASVs by prokaryotic cell counts obtained from flow cytometry analysis. NMDS plots were generated by calculating Bray-Curtis dissimilarity scores based on ASV relative abundances using the *vegdist()* function in R.

### Self-Organizing Map

The Kohonen package (32) in R (v. 4.0.0) was used to cluster samples based on microbial community composition. A self-organizing map was constructed using ASV relative abundances and each sample was assigned to a map unit based on community similarities. Our map was composed of 4x4 circular nodes in a hexagonal, nontoroidal configuration. Clusters were generated using k-means clustering and a reasonable value for k was chosen by estimating the point of inflection in a scree plot of within-clusters sum of squares (Figure S6).

### Discriminative taxa identification

Generalized joint attribute modelling (GJAM) was applied to model the 250 most abundant ASVs and select environmental factors (temperature, salinity, chlorophyll *a*, PO_4_), using the GJAM v. 2.6.2 package in R. Iteration was set at 20,000 and burning at 5,000. Results were visualized using the built-in function ‘gjamPlot’. In order to identify discriminative taxa between SOM clusters, the 250 most abundant ASVs were analyzed with LEfSe (41). The threshold for significance on the logarithmic LDA score for discriminative features was two. P-values were obtained from the Kruskall-Wallis test for differences in ASV abundances among SOM cluster assignments. These two metrics differ in approach: LEfSe identifies phylotype enrichment in a specific cluster while GJAM can identify the magnitude and directionality (positive or negative) of associations across multiple clusters. ASVs exhibiting differential abundances between the Gulf Stream and eddy were further identified using DESeq2 (42). Only ASVs with an average relative abundance >0.05% averaged across samples being compared (Gulf Stream and eddy) were included. Significant differences were considered those with a Benjamin-Hochberg multiple hypothesis adjusted p<0.05. Quantification of the processes shaping microbiome community assembly using Beta Mean Nearest Taxon Distance (βMNTD) (5) was not possible because our data did not exhibit the required phylogenetic signal (non-significant Mantel correlogram).

### Satellite Data

Satellite data from two sources is used in this work. Chlorophyll *a* products were derived from ocean color imagery acquired by the European Commission Copernicus programme’s Sentinel-3 Ocean and Land Color Instrument (OLCI) using the CHL_OC4ME blue to green band ratio algorithm (62). The OLCI data was used to track surface chlorophyll *a* within the eddy over its lifetime. Sea surface temperature products were from the Geostationary Operational Environmental Satellite 16 (GOES-16) which uses a retrieval from the long wave infrared bands (8μm-12μm) (63). OLCI Level 2 and obtained from https://coda.eumetsat.int and GOES-16 hourly SST was acquired from https://cwcgom.aoml.noaa.gov/erddap/griddap/goes16SSThourly.html.

### Physical Water Parcel Delineations

Temperature and salinity delineations between water parcels are specific to this region and time. They are based on satellite tracking of the eddy water mass and known Temperature-Salinity properties of these waters. In general Gulf Stream waters are the hottest and saltiest, South Atlantic Bight waters are slightly fresher and cooler, Mid Atlantic Bight slope waters are cooler and fresher again, and Mid Atlantic Bight shelf water is the coolest and freshest (31, 64). To determine the exact thresholds of temperature and salinity these general patterns were informed and reinforced by observed sharp frontal gradients between water masses. This led to the following thresholds: Gulf Stream > 28.2 °C, eddy <28.2 °C and > 35.75 PSU, continental slope <35.75 PSU and > 34 PSU, and continental shelf <34 PSU (Figure S1) (31).

### Data and Code Repository

Cruise field data is deposited: <u>doi.org/10.5281/zenodo.7680135</u> and SSU rRNA library sequences are deposited as part of Bioproject PRJNA868376. Analysis scripts are available at https://github.com/jgronniger/Gulf-Stream-Eddy-2021.

## Acknowledgements

We acknowledge the assistance of the captain and crew of the *R/V Shearwater*.

## CRediT Contributions

**Jessica Gronniger-** Conceptualization; Data Curation; Formal Analysis; Visualization; Writing-original draft

**Patrick C. Gray-** Conceptualization; Data Curation; Formal Analysis; Visualization; Writing-review and editing

**Alexandria K. Niebergall-** Formal Analysis; Writing-review and editing

**Zackary I. Johnson-** Resources; Writing-review and editing

**Dana E. Hunt-** Conceptualization; Resources; Writing-original draft;

## Supplemental materials

Table S1. Multiple regression analysis of select environmental variables on Bray-Curtis dissimilarity for 16S rRNA gene libraries using ‘envfit’ in R (v 4.0.0) with 9999 permutations.

Table S2. Linear discriminant analysis effect size (LEfSE) results for the 250 most abundant ASVs across all samples.

Table S3. GJAM beta-coefficients of ASVs found to have a significant association with at least one SOM cluster.

Table S4. Linear discriminant analysis effect size (LEfSE) results for the 250 most abundant ASVs in Gulf Stream and eddy comparison.

Figure S1. Environmental variables for physical parameter defined water parcels (A) Temperature and (B) salinity for each of the temperature and salinity-defined water parcels (Continental Shelf, Continental Slope, Gulf Stream, Eddy) as determined using the flow-through seawater system at ∼1m depth. Center lines denotes the median value while the box encompasses the 25^th^ to 75^th^ percentiles of the dataset. Whiskers denote the 5^th^ and 95^th^ percentiles and values that fall outside these ranges are shown as individual points. Brackets indicate p-values associated with Wilcoxon Rank Sum test pairwise comparisons.

Figure S2. Non-metric multidimensional scaling (NMDS) ordination comparisons of bacterial communities grouped by physical and biological parameters. NMDS computed based on Bray-Curtis dissimilarity for 16S rRNA gene libraries and colored by (A) oceanographic water parcels (temperature and salinity) and (B) Self-Organizing Map (SOM) clusters based on the microbiome. Ellipses show the multivariate t-distribution 95% confidence interval for the mean microbial community of each water parcel or SOM cluster.

Figure S3. Maps of sampling sites for five transects off the coast of Cape Hatteras, North Carolina, USA from September 5-7, 2021. Background color represents sea surface temperatures at time of sampling obtained from the Geostationary Operational Environmental Satellite 16 (GOES-16). Each sampling point is shown along the transects and categorized by oceanographic water parcel (shape), defined by temperature and salinity, and microbial-community based k-means clusters (color), generated using the self-organizing map R package ‘kohonen’. Black dotted lines indicate the approximate outline of the frontal eddy during the time of sampling based on a qualitative analysis of the SST imagery.

Figure S4. Environmental conditions for Self-Organizing Map (SOM) clusters. Numbered clusters correspond to the most common environments: Continental shelf (Cluster 1), Continental slope (Cluster 2), Gulf Stream (Cluster 3), Eddy (Cluster 4). For the box and whisker plots, the center lines denote the median value while the box encompasses the 25^th^ to 75^th^ percentiles of the dataset. Whiskers denote the 5^th^ and 95^th^ percentiles and values that fall outside these ranges are shown as individual points. Brackets indicate p-values of Wilcoxon Rank Sum pairwise comparisons.

Figure S5. Non-metric multidimensional scaling (NMDS) ordination computed based on Sørensen similarity for 16S rRNA gene libraries. Each sampling point is categorized by oceanographic water parcel (shapes), defined by temperature and salinity, and microbial-community based k-means clusters (colors), generated using the self-organizing map R package ‘kohonen’. Ellipses show the multivariate t-distribution 95% confidence interval for the mean of each Self-Organizing Map (SOM cluster).

Figure S6. Self-Organizing Map within-clusters sum of squares plot for cluster number selection. Calculated using a 4x4 grid with hexagonal mapping units in the ‘kohonen’ (v.3.0.11) in R (v4.0.0).

