## Supplemental Figures for "A Gulf Stream frontal eddy harbors a distinct microbiome compared to adjacent waters"

**Table S1. Multiple regression analysis of select environmental variables on Bray-Curtis dissimilarity for 16S rRNA gene libraries using 'envfit' in R (v 4.0.0) with 9999 permutations.**

| Environmental Parameter | $R^2$ | p-value[1] |
|---|---|---|
| Temperature | 0.7517 | **0.0011*** |
| Salinity | 0.8704 | **0.0011*** |
| *Synechococcus* | 0.8934 | **0.0011*** |
| Picocyanobacteria | 0.3373 | **0.0055*** |
| Bacteria | 0.8067 | **0.0011*** |
| Chlorophyll *a* | 0.5448 | **0.0011*** |
| $NO_3$ | 0.1165 | 0.8173 |
| $PO_4$ | 0.4381 | **0.0011*** |
| $SiO_4$ | 0.1976 | 0.1133 |
| $NH_4$ | 0.0778 | 1.0000 |
| $NO_2$ | 0.0059 | 1.0000 |

[1] Bonferroni-corrected p-values, those below the significance threshold (p<0.05) are indicated by bold and an '*'.

**Table S2. Linear discriminant analysis effect size (LEfSE) results for the 250 most abundant ASVs across all samples.**

| SOM Cluster | ASV[1] | LDA Score | RDP Classifier Taxonomy[2] |
|---|---|---|---|
| Shelf (1) | 15324 | 3.73328408 | k__Bacteria; p__Cyanobacteria; c__Synechococcophycideae; o__Synechococcales; f__Synechococcaceae; g__Synechococcus; s__ |
| | 15364 | 4.9590123 | k__Bacteria; p__Cyanobacteria; c__Synechococcophycideae; o__Synechococcales; f__Synechococcaceae; g__Synechococcus; s__ |
| | 16066 | 3.78628889 | k__Bacteria; p__Proteobacteria; c__Alphaproteobacteria; o__Rickettsiales; f__Pelagibacteraceae; g__; s__ |
| | 25676 | 3.93839911 | k__Bacteria; p__Proteobacteria; c__Alphaproteobacteria; o__Rickettsiales; f__Pelagibacteraceae; g__; s__ |
| | 25688 | 3.98091799 | k__Bacteria; p__Proteobacteria; c__Alphaproteobacteria; o__Rickettsiales; f__Pelagibacteraceae; g__; s__ |
| | 28783 | 3.80741984 | k__Bacteria; p__Cyanobacteria; c__Synechococcophycideae; o__Synechococcales; f__Synechococcaceae; g__Synechococcus; s__ |
| | 28838 | 3.89548198 | k__Bacteria; p__Cyanobacteria; c__Synechococcophycideae; o__Synechococcales; f__Synechococcaceae; g__Synechococcus; s__ |
| | 29391 | 3.6573079 | k__Bacteria; p__Proteobacteria; c__Alphaproteobacteria; o__Rickettsiales; f__Pelagibacteraceae; g__; s__ |
| | 29415 | 3.85572743 | k__Bacteria; p__Proteobacteria; c__Alphaproteobacteria; o__Rickettsiales; f__Pelagibacteraceae; g__; s__ |
| | 33047 | 3.60957607 | k__Bacteria; p__Planctomycetes; c__Planctomycetia; o__Pirellulales; f__Pirellulaceae |
| | 37782 | 3.72072291 | k__Bacteria; p__Cyanobacteria; c__Synechococcophycideae; o__Synechococcales; f__Synechococcaceae; g__Synechococcus; s__ |
| | 37872 | 3.75027342 | k__Bacteria; p__Proteobacteria; c__Alphaproteobacteria; o__Rickettsiales; f__Pelagibacteraceae; g__; s__ |
| | 37887 | 4.49670087 | k__Bacteria; p__Proteobacteria; c__Alphaproteobacteria; o__Rickettsiales; f__Pelagibacteraceae; g__; s__ |
| | 37894 | 3.95154144 | k__Bacteria; p__Proteobacteria; c__Alphaproteobacteria; o__Rickettsiales; f__Pelagibacteraceae; g__; s__ |
| | 39899 | 3.75740656 | k__Bacteria; p__Actinobacteria; c__Acidimicrobiia; o__Acidimicrobiales; f__OCS155; g__; s__ |
| | 40663 | 3.91135983 | k__Bacteria; p__Proteobacteria; c__Alphaproteobacteria; o__Rickettsiales; f__Pelagibacteraceae; g__; s__ |
| | 41353 | 3.6671388 | k__Bacteria; p__Proteobacteria; c__Alphaproteobacteria; o__; f__; g__; s__ |
| Slope (2) | 29355 | 3.67470643 | k__Bacteria; p__Proteobacteria; c__Alphaproteobacteria; o__Rickettsiales; f__Pelagibacteraceae; g__; s__ |
| | 38421 | 3.58789326 | k__Bacteria; p__Proteobacteria; c__Alphaproteobacteria; o__Rickettsiales; f__Pelagibacteraceae; g__; s__ |
| | 40988 | 3.76799534 | k__Archaea; p__Euryarchaeota; c__Thermoplasmata; o__E2; f__Marine group II; g__; s__ |
| Gulf Stream (3) | 11073 | 3.90423733 | k__Bacteria; p__SBR1093; c__A712011; o__; f__; g__; s__ |

| | | | |
|---|---|---|---|
| Gulf Stream (3) | 16069 | 3.6882494 | k__Bacteria; p__Proteobacteria; c__Alphaproteobacteria; o__Rickettsiales; f__Pelagibacteraceae; g__; s__ |
| | 27661 | 3.81946893 | k__Bacteria; p__Proteobacteria; c__Alphaproteobacteria; o__Rickettsiales; f__Pelagibacteraceae; g__; s__ |
| | 30202 | 3.56668482 | k__Bacteria; p__Proteobacteria; c__Alphaproteobacteria; o__Rickettsiales; f__Pelagibacteraceae; g__; s__ |
| | 32746 | 3.46196378 | k__Bacteria; p__Bacteroidetes; c__Cytophagia; o__Cytophagales; f__Flammeovirgaceae; g__; s__ |
| | 34223 | 4.22025463 | k__Bacteria; p__Actinobacteria; c__Acidimicrobiia; o__Acidimicrobiales; f__OCS155; g__; s__ |
| | 35277 | 3.95874603 | k__Bacteria; p__Proteobacteria; c__Gammaproteobacteria; o__HTCC2188; f__HTCC2089; g__; s__ |
| | 36431 | 3.54826415 | k__Bacteria; p__Proteobacteria; c__Alphaproteobacteria; o__Rickettsiales; f__AEGEAN_112; g__; s__ |
| | 36482 | 3.77150061 | k__Bacteria; p__Proteobacteria; c__Alphaproteobacteria; o__Rickettsiales; f__AEGEAN_112; g__; s__ |
| | 37115 | 3.60940897 | k__Bacteria; p__Proteobacteria; c__Alphaproteobacteria; o__Rhodobacterales; f__Rhodobacteraceae; g__; s__ |
| | 38991 | 3.4652386 | k__Bacteria; p__Proteobacteria; c__Alphaproteobacteria; o__; f__; g__; s__ |
| | 40444 | 3.63206755 | k__Bacteria; p__Bacteroidetes; c__Flavobacteriia; o__Flavobacteriales; f__Flavobacteriaceae; g__; s__ |
| | 40517 | 3.70144398 | k__Bacteria; p__Proteobacteria; c__Alphaproteobacteria; o__Rickettsiales; f__Pelagibacteraceae; g__; s__ |
| | 41180 | 3.3663844 | k__Bacteria; p__Proteobacteria; c__Alphaproteobacteria; o__; f__; g__; s__ |
| Eddy (4) | 17637 | 5.19192744 | k__Bacteria; p__Cyanobacteria; c__Synechococcophycideae; o__Synechococcales; f__Synechococcaceae; g__Prochlorococcus; s__ |
| | 25757 | 3.85606733 | k__Bacteria; p__Actinobacteria; c__Acidimicrobiia; o__Acidimicrobiales; f__OCS155; g__; s__ |
| | 31071 | 4.09113514 | k__Bacteria; p__Cyanobacteria; c__Synechococcophycideae; o__Synechococcales; f__Synechococcaceae; g__Prochlorococcus; s__ |
| | 31171 | 3.78454931 | k__Bacteria; p__Cyanobacteria; c__Synechococcophycideae; o__Synechococcales; f__Synechococcaceae; g__Prochlorococcus; s__ |
| | 31582 | 3.98447696 | k__Bacteria; p__Cyanobacteria; c__Synechococcophycideae; o__Synechococcales; f__Synechococcaceae; g__Prochlorococcus; s__ |
| | 40886 | 3.69902041 | k__Bacteria; p__Cyanobacteria; c__Synechococcophycideae; o__Synechococcales; f__Synechococcaceae; g__Prochlorococcus; s__ |

[1]Only significant discriminative taxa ($p < 0.05$) and their respective linear discriminant analysis (LDA) scores are reported for each SOM cluster.
[2]Taxonomies were assigned based on RDP classifier.

**Table S3. GJAM beta-coefficients of ASVs found to have a significant association with at least one SOM cluster.**

| ASV | Shelf (1) | Slope (2) | Gulf Stream (3) | Eddy (4) | RDP Classifier Taxonomy[1] |
|---|---|---|---|---|---|
| 26179 | **0.45**[2] | 0.0925 | -0.264 | -0.279 | k__Bacteria: p__Cyanobacteria: c__Chloroplast: o__Stramenopiles: f__: g__: s__ |
| 34223 | **-1.35** | **0.0971** | **0.767** | **0.489** | k__Bacteria: p__Proteobacteria: c__Alphaproteobacteria: o__Rickettsiales: f__Pelagibacteraceae: g__: s__ |
| 39899 | **0.794** | 0.429 | **-0.554** | **-0.669** | k__Bacteria: p__Cyanobacteria: c__Chloroplast: o__Stramenopiles: f__: g__: s__ |
| 39875 | **2.68** | **0.805** | **-1.72** | **-1.76** | k__Bacteria: p__Verrucomicrobia: c__Opitutae: o__Puniceicoccales: f__Puniceicoccaceae: g__Coraliomargarita: s__ |
| 15364 | **0.00592** | 0.00136 | **-0.00364** | **-0.00364** | k__Bacteria: p__Cyanobacteria: c__Chloroplast: o__Stramenopiles: f__: g__: s__ |
| 31461 | **0.19** | 0.413 | -0.296 | -0.307 | k__Bacteria: p__Proteobacteria: c__Alphaproteobacteria: o__Rickettsiales: f__Pelagibacteraceae: g__: s__ |
| 31501 | **0.0709** | 0.471 | -0.29 | -0.252 | k__Bacteria: p__Cyanobacteria: c__Chloroplast: o__Stramenopiles: f__: g__: s__ |
| 17637 | **-0.00586** | -0.00104 | **0.00321** | **0.00369** | k__Bacteria; p__Cyanobacteria; c__Synechococcophycideae; o__Synechococcales; f__Synechococcaceae; g__Prochlorococcus; s__ |
| 30907 | **-1.92** | **0.41** | **-0.44** | **1.95** | k__Bacteria; p__Actinobacteria; c__Acidimicrobiia; o__Acidimicrobiales; f__OCS155; g__; s__ |
| 14960 | **-0.182** | 0.28 | -0.115 | 0.0167 | k__Bacteria; p__Cyanobacteria; c__Synechococcophycideae; o__Synechococcales; f__Synechococcaceae |
| 28680 | **0.0178** | 0.323 | -0.191 | -0.149 | k__Bacteria; p__Cyanobacteria; c__Synechococcophycideae; o__Synechococcales; f__Synechococcaceae; g__Prochlorococcus; s__ |
| 41180 | **0.398** | -0.0919 | -0.103 | -0.204 | k__Bacteria; p__Cyanobacteria; c__Synechococcophycideae; o__Synechococcales; f__Synechococcaceae; g__Synechococcus; s__ |
| 41241 | **0.775** | -0.0334 | **-0.261** | **-0.481** | k__Bacteria; p__Cyanobacteria; c__Synechococcophycideae; o__Synechococcales; f__Synechococcaceae |
| 40517 | **1.16** | 0.294 | **-0.456** | **-0.997** | k__Bacteria; p__Proteobacteria; c__Alphaproteobacteria; o__; f__; g__; s__ |
| 40560 | **0.135** | 0.138 | -0.0614 | -0.211 | k__Bacteria; p__Proteobacteria; c__Alphaproteobacteria; o__Rickettsiales; f__Pelagibacteraceae; g__; s__ |
| 37271 | **0.378** | 0.0203 | -0.16 | -0.238 | k__Bacteria; p__Proteobacteria; c__Alphaproteobacteria; o__Rickettsiales; f__Pelagibacteraceae; g__; s__ |

| ASV | Shelf (1) | Slope (2) | Gulf Stream (3) | Eddy (4) | RDP Classifier Taxonomy[1] |
|---|---|---|---|---|---|
| 41364 | **0.158** | 0.332 | -0.22 | -0.27 | k__Bacteria; p__Cyanobacteria; c__Synechococcophycideae; o__Synechococcales; f__Synechococcaceae |
| 40663 | **0.664** | 0.148 | **-0.362** | **-0.449** | k__Bacteria; p__Proteobacteria; c__Alphaproteobacteria; o__Rickettsiales; f__Pelagibacteraceae; g__; s__ |
| 37887 | **5.38** | **1.27** | **-3** | **-3.65** | k__Bacteria; p__Proteobacteria; c__Gammaproteobacteria; o__Oceanospirillales; f__Halomonadaceae; g__Candidatus Portiera; s__ |
| 38437 | **3.28** | **2.2** | **-2.66** | **-2.82** | k__Bacteria; p__Cyanobacteria; c__Synechococcophycideae; o__Synechococcales; f__Synechococcaceae; g__Synechococcus; s__ |
| 30202 | **0.743** | 0.249 | **-0.294** | **-0.697** | k__Bacteria; p__Proteobacteria; c__Alphaproteobacteria; o__Rickettsiales; f__Pelagibacteraceae; g__; s__ |
| 25676 | **1.16** | **-0.0622** | **-0.468** | **-0.625** | k__Bacteria; p__Proteobacteria; c__Alphaproteobacteria; o__; f__; g__; s__ |

[1] Taxonomies were assigned by RDP classifier.

[2] Bolded values indicated the 95% CI of the Bayesian factor does not include 0.

**Table S4. Linear discriminant analysis effect size (LEfSE) results for the 250 most abundant ASVs in Gulf Stream and eddy comparison.**

| SOM Cluster | ASV[1] | LDA Score | RDP Classifier Taxonomy[2] |
|---|---|---|---|
| Gulf Stream (3) | 41241 | 3.18594313 | k__Bacteria p__Proteobacteria c__Alphaproteobacteria o__ f__ g__ s__ |
| | 35723 | 2.87802661 | k__Bacteria p__Proteobacteria c__Alphaproteobacteria o__ f__ g__ s__ |
| | 35786 | 2.64468697 | k__Bacteria p__Proteobacteria c__Alphaproteobacteria o__ f__ g__ s__ |
| | 41259 | 3.41917204 | k__Bacteria p__Proteobacteria c__Alphaproteobacteria o__ f__ g__ s__ |
| | 18612 | 2.72335095 | k__Bacteria p__Proteobacteria c__Alphaproteobacteria o__ f__ g__ s__ |
| | 41253 | 2.8564117 | k__Bacteria p__Proteobacteria c__Alphaproteobacteria o__ f__ g__ s__ |
| | 40306 | 2.81500575 | k__Bacteria p__Proteobacteria c__Alphaproteobacteria o__ f__ g__ s__ |
| | 33721 | 2.68013423 | k__Bacteria p__Cyanobacteria c__Chloroplast o__Stramenopiles f__ g__ s__ |
| | 38991 | 2.77872419 | k__Bacteria p__Proteobacteria c__Alphaproteobacteria o__ f__ g__ s__ |
| | 30093 | 2.43129345 | k__Bacteria p__Proteobacteria c__Alphaproteobacteria o__ f__ g__ s__ |
| | 39026 | 2.64906179 | k__Bacteria p__Proteobacteria c__Alphaproteobacteria o__ f__ g__ s__ |
| | 41197 | 2.61240735 | k__Bacteria p__Proteobacteria c__Alphaproteobacteria o__ f__ g__ s__ |
| | 39700 | 2.6341518 | k__Bacteria p__Proteobacteria c__Alphaproteobacteria o__ f__ g__ s__ |
| | 41353 | 2.71014553 | k__Bacteria p__Proteobacteria c__Alphaproteobacteria o__ f__ g__ s__ |
| | 41180 | 3.04937441 | k__Bacteria p__Proteobacteria c__Alphaproteobacteria o__ f__ g__ s__ |
| | 26228 | 2.65937611 | k__Bacteria p__Proteobacteria c__Alphaproteobacteria o__Rickettsiales f__AEGEAN_112 g__ s__ |
| | 36402 | 2.65863134 | k__Bacteria p__Proteobacteria c__Alphaproteobacteria o__Rickettsiales f__AEGEAN_112 g__ s__ |
| | 40361 | 2.6412984 | k__Bacteria p__Proteobacteria c__Alphaproteobacteria o__Rickettsiales f__AEGEAN_112 g__ s__ |
| | 36431 | 2.67394529 | k__Bacteria p__Proteobacteria c__Alphaproteobacteria o__Rickettsiales f__AEGEAN_112 g__ s__ |
| | 32746 | 2.72709689 | k__Bacteria p__Bacteroidetes c__Cytophagia o__Cytophagales f__Flammeovirgaceae g__ s__ |
| | 40423 | 2.77694525 | k__Bacteria p__Bacteroidetes c__Flavobacteriia o__Flavobacteriales f__Flavobacteriaceae g__ s__ |

| SOM Cluster | ASV[1] | LDA Score | RDP Classifier Taxonomy[2] |
|---|---|---|---|
| Gulf Stream (3) | 36891 | 2.75734527 | k__Bacteria p__Bacteroidetes c__Flavobacteriia o__Flavobacteriales f__Flavobacteriaceae g__ s__ |
| | 40454 | 2.58415269 | k__Bacteria p__Bacteroidetes c__Flavobacteriia o__Flavobacteriales f__Flavobacteriaceae g__ s__ |
| | 40444 | 2.68321664 | k__Bacteria p__Bacteroidetes c__Flavobacteriia o__Flavobacteriales f__Flavobacteriaceae g__ s__ |
| | 40184 | 2.72751219 | k__Bacteria p__Proteobacteria c__Gammaproteobacteria o__Oceanospirillales f__Halomonadaceae g__Candidatus Portiera s__ |
| | 36269 | 2.62150988 | k__Bacteria p__Proteobacteria c__Gammaproteobacteria o__Oceanospirillales f__Halomonadaceae g__Candidatus Portiera s__ |
| | 35277 | 2.54453919 | k__Bacteria p__Proteobacteria c__Gammaproteobacteria o__HTCC2188 f__HTCC2089 g__ s__ |
| | 39899 | 3.03396249 | k__Bacteria p__Actinobacteria c__Acidimicrobiia o__Acidimicrobiales f__OCS155 g__ s__ |
| | 22760 | 2.68076051 | k__Bacteria p__Actinobacteria c__Acidimicrobiia o__Acidimicrobiales f__OCS155 g__ s__ |
| | 34223 | 3.57733799 | k__Bacteria p__Actinobacteria c__Acidimicrobiia o__Acidimicrobiales f__OCS155 g__ s__ |
| | 37435 | 3.29699046 | k__Bacteria p__Proteobacteria c__Alphaproteobacteria o__Rickettsiales f__Pelagibacteraceae g__ s__ |
| | 16069 | 2.75373557 | k__Bacteria p__Proteobacteria c__Alphaproteobacteria o__Rickettsiales f__Pelagibacteraceae g__ s__ |
| | 37296 | 2.92371969 | k__Bacteria p__Proteobacteria c__Alphaproteobacteria o__Rickettsiales f__Pelagibacteraceae g__ s__ |
| | 38073 | 2.64281315 | k__Bacteria p__Proteobacteria c__Alphaproteobacteria o__Rickettsiales f__Pelagibacteraceae g__ s__ |
| | 37887 | 3.7428403 | k__Bacteria p__Proteobacteria c__Alphaproteobacteria o__Rickettsiales f__Pelagibacteraceae g__ s__ |
| | 40671 | 2.82917305 | k__Bacteria p__Proteobacteria c__Alphaproteobacteria o__Rickettsiales f__Pelagibacteraceae g__ s__ |
| | 30202 | 3.51842188 | k__Bacteria p__Proteobacteria c__Alphaproteobacteria o__Rickettsiales f__Pelagibacteraceae g__ s__ |
| | 29635 | 3.17568233 | k__Bacteria p__Proteobacteria c__Alphaproteobacteria o__Rickettsiales f__Pelagibacteraceae g__ s__ |
| | 40686 | 3.30805217 | k__Bacteria p__Proteobacteria c__Alphaproteobacteria o__Rickettsiales f__Pelagibacteraceae g__ s__ |
| | 37438 | 2.89580175 | k__Bacteria p__Proteobacteria c__Alphaproteobacteria o__Rickettsiales f__Pelagibacteraceae g__ s__ |
| | 40517 | 3.69185831 | k__Bacteria p__Proteobacteria c__Alphaproteobacteria o__Rickettsiales f__Pelagibacteraceae g__ s__ |
| | 27661 | 2.6098161 | k__Bacteria p__Proteobacteria c__Alphaproteobacteria o__Rickettsiales f__Pelagibacteraceae g__ s__ |
| | 29562 | 3.03800976 | k__Bacteria p__Proteobacteria c__Alphaproteobacteria o__Rickettsiales f__Pelagibacteraceae g__ s__ |
| | 37334 | 2.72094002 | k__Bacteria p__Proteobacteria c__Alphaproteobacteria o__Rickettsiales f__Pelagibacteraceae g__ s__ |

| SOM Cluster | ASV[1] | LDA Score | RDP Classifier Taxonomy[2] |
|---|---|---|---|
| Gulf Stream (3) | 37427 | 2.83161429 | k__Bacteria p__Proteobacteria c__Alphaproteobacteria o__Rickettsiales f__Pelagibacteraceae g__ s__ |
| | 38028 | 2.75159287 | k__Bacteria p__Proteobacteria c__Alphaproteobacteria o__Rickettsiales f__Pelagibacteraceae g__ s__ |
| | 29511 | 2.55897771 | k__Bacteria p__Proteobacteria c__Alphaproteobacteria o__Rickettsiales f__Pelagibacteraceae g__ s__ |
| | 40663 | 2.71380706 | k__Bacteria p__Proteobacteria c__Alphaproteobacteria o__Rickettsiales f__Pelagibacteraceae g__ s__ |
| | 37401 | 2.86527135 | k__Bacteria p__Proteobacteria c__Alphaproteobacteria o__Rickettsiales f__Pelagibacteraceae g__ s__ |
| | 29523 | 2.6283924 | k__Bacteria p__Proteobacteria c__Alphaproteobacteria o__Rickettsiales f__Pelagibacteraceae g__ s__ |
| | 40560 | 3.13838252 | k__Bacteria p__Proteobacteria c__Alphaproteobacteria o__Rickettsiales f__Pelagibacteraceae g__ s__ |
| | 27672 | 2.66652697 | k__Bacteria p__Proteobacteria c__Alphaproteobacteria o__Rickettsiales f__Pelagibacteraceae g__ s__ |
| | 29391 | 2.64675611 | k__Bacteria p__Proteobacteria c__Alphaproteobacteria o__Rickettsiales f__Pelagibacteraceae g__ s__ |
| | 40779 | 2.63742589 | k__Bacteria p__Proteobacteria c__Alphaproteobacteria o__Rickettsiales f__Pelagibacteraceae g__ s__ |
| | 27803 | 2.716869 | k__Bacteria p__Proteobacteria c__Alphaproteobacteria o__Rickettsiales f__Pelagibacteraceae g__ s__ |
| | 30030 | 2.54515922 | k__Bacteria p__Proteobacteria c__Alphaproteobacteria o__Rickettsiales f__Pelagibacteraceae g__ s__ |
| | 37872 | 2.60447871 | k__Bacteria p__Proteobacteria c__Alphaproteobacteria o__Rickettsiales f__Pelagibacteraceae g__ s__ |
| | 29653 | 2.6433467 | k__Bacteria p__Proteobacteria c__Alphaproteobacteria o__Rickettsiales f__Pelagibacteraceae g__ s__ |
| | 37325 | 2.37617713 | k__Bacteria p__Proteobacteria c__Alphaproteobacteria o__Rickettsiales f__Pelagibacteraceae g__ s__ |
| | 38278 | 2.68376136 | k__Bacteria p__Proteobacteria c__Alphaproteobacteria o__Rickettsiales f__Pelagibacteraceae g__ s__ |
| | 30044 | 2.33264077 | k__Bacteria p__Proteobacteria c__Alphaproteobacteria o__Rickettsiales f__Pelagibacteraceae g__ s__ |
| | 34183 | 2.58178787 | k__Bacteria p__Proteobacteria c__Alphaproteobacteria o__Rickettsiales f__Pelagibacteraceae g__ s__ |
| | 38086 | 2.63638699 | k__Bacteria p__Proteobacteria c__Alphaproteobacteria o__Rickettsiales f__Pelagibacteraceae g__ s__ |
| | 29508 | 2.6264426 | k__Bacteria p__Proteobacteria c__Alphaproteobacteria o__Rickettsiales f__Pelagibacteraceae g__ s__ |
| | 29371 | 2.602273 | k__Bacteria p__Proteobacteria c__Alphaproteobacteria o__Rickettsiales f__Pelagibacteraceae g__ s__ |
| | 38374 | 2.59349408 | k__Bacteria p__Proteobacteria c__Alphaproteobacteria o__Rickettsiales f__Pelagibacteraceae g__ s__ |
| | 40552 | 2.51892763 | k__Bacteria p__Proteobacteria c__Alphaproteobacteria o__Rickettsiales f__Pelagibacteraceae g__ s__ |

| SOM Cluster | ASV[1] | LDA Score | RDP Classifier Taxonomy[2] |
|---|---|---|---|
| Gulf Stream (3) | 41297 | 3.25845343 | k__Bacteria p__Cyanobacteria c__Oscillatoriophycideae o__Oscillatoriales f__Phormidiaceae g__Trichodesmium s__ |
| | 37099 | 2.51909067 | k__Bacteria p__Proteobacteria c__Alphaproteobacteria o__Rhodobacterales f__Rhodobacteraceae g__ s__ |
| | 29023 | 2.8648328 | k__Bacteria p__Cyanobacteria c__Synechococcophycideae o__Synechococcales f__Synechococcaceae g__Synechococcus s__ |
| | 15436 | 2.97703905 | k__Bacteria p__Cyanobacteria c__Synechococcophycideae o__Synechococcales f__Synechococcaceae g__Synechococcus s__ |
| | 28989 | 2.89043595 | k__Bacteria p__Cyanobacteria c__Synechococcophycideae o__Synechococcales f__Synechococcaceae g__Synechococcus s__ |
| | 37718 | 2.5864231 | k__Bacteria p__Cyanobacteria c__Synechococcophycideae o__Synechococcales f__Synechococcaceae g__Synechococcus s__ |
| | 15364 | 4.00676486 | k__Bacteria p__Cyanobacteria c__Synechococcophycideae o__Synechococcales f__Synechococcaceae g__Synechococcus s__ |
| | 31430 | 2.48657345 | k__Bacteria p__Cyanobacteria c__Synechococcophycideae o__Synechococcales f__Synechococcaceae |
| | 26590 | 2.73118048 | k__Bacteria p__Proteobacteria c__Alphaproteobacteria |
| | 40393 | 2.72359348 | k__Bacteria p__Proteobacteria c__Alphaproteobacteria |
| | 36801 | 2.60200759 | k__Bacteria p__Proteobacteria c__Alphaproteobacteria |
| Eddy (4) | 37527 | 2.60111642 | k__Bacteria p__SAR406 c__AB16 o__Arctic96B-7 f__A714017 g__SGSH944 s__ |
| | 37589 | 2.54754652 | k__Bacteria p__SAR406 c__AB16 o__Arctic96B-7 f__A714017 g__SGSH944 s__ |
| | 40604 | 2.68067103 | k__Bacteria p__SAR406 c__AB16 o__Arctic96B-7 f__A714017 g__SGSH944 s__ |
| | 25757 | 2.69005479 | k__Bacteria p__Actinobacteria c__Acidimicrobiia o__Acidimicrobiales f__OCS155 g__ s__ |
| | 17637 | 4.76169019 | k__Bacteria p__Cyanobacteria c__Synechococcophycideae o__Synechococcales f__Synechococcaceae g__Prochlorococcus s__ |
| | 31171 | 3.26373153 | k__Bacteria p__Cyanobacteria c__Synechococcophycideae o__Synechococcales f__Synechococcaceae g__Prochlorococcus s__ |
| | 30907 | 4.0984162 | k__Bacteria p__Cyanobacteria c__Synechococcophycideae o__Synechococcales f__Synechococcaceae g__Prochlorococcus s__ |
| | 18179 | 2.88142993 | k__Bacteria p__Cyanobacteria c__Synechococcophycideae o__Synechococcales f__Synechococcaceae g__Prochlorococcus s__ |
| | 30866 | 2.73090441 | k__Bacteria p__Cyanobacteria c__Synechococcophycideae o__Synechococcales f__Synechococcaceae g__Prochlorococcus s__ |
| | 31582 | 2.85593135 | k__Bacteria p__Cyanobacteria c__Synechococcophycideae o__Synechococcales f__Synechococcaceae g__Prochlorococcus s__ |
| | 40886 | 2.8132281 | k__Bacteria p__Cyanobacteria c__Synechococcophycideae o__Synechococcales f__Synechococcaceae g__Prochlorococcus s__ |
| | 30962 | 2.70538611 | k__Bacteria p__Cyanobacteria c__Synechococcophycideae o__Synechococcales f__Synechococcaceae g__Prochlorococcus s__ |

| SOM Cluster | ASV[1] | LDA Score | RDP Classifier Taxonomy[2] |
|---|---|---|---|
| Eddy (4) | 18024 | 2.60242015 | k__Bacteria p__Cyanobacteria c__Synechococcophycideae o__Synechococcales f__Synechococcaceae g__Prochlorococcus s__ |
| | 31265 | 2.63181595 | k__Bacteria p__Cyanobacteria c__Synechococcophycideae o__Synechococcales f__Synechococcaceae g__Prochlorococcus s__ |
| | 31645 | 2.71037166 | k__Bacteria p__Cyanobacteria c__Synechococcophycideae o__Synechococcales f__Synechococcaceae |
| | 31071 | 2.62434453 | k__Bacteria p__Cyanobacteria c__Synechococcophycideae o__Synechococcales f__Synechococcaceae g__Prochlorococcus s__ |
| | 38742 | 2.57431276 | k__Bacteria p__Cyanobacteria c__Synechococcophycideae o__Synechococcales f__Synechococcaceae g__Prochlorococcus s__ |

[1]Only significant discriminative taxa ($p < 0.05$) and their respective linear discriminant analysis (LDA) scores are reported for each SOM cluster.
[2]Taxonomies were assigned based on RDP classifier.

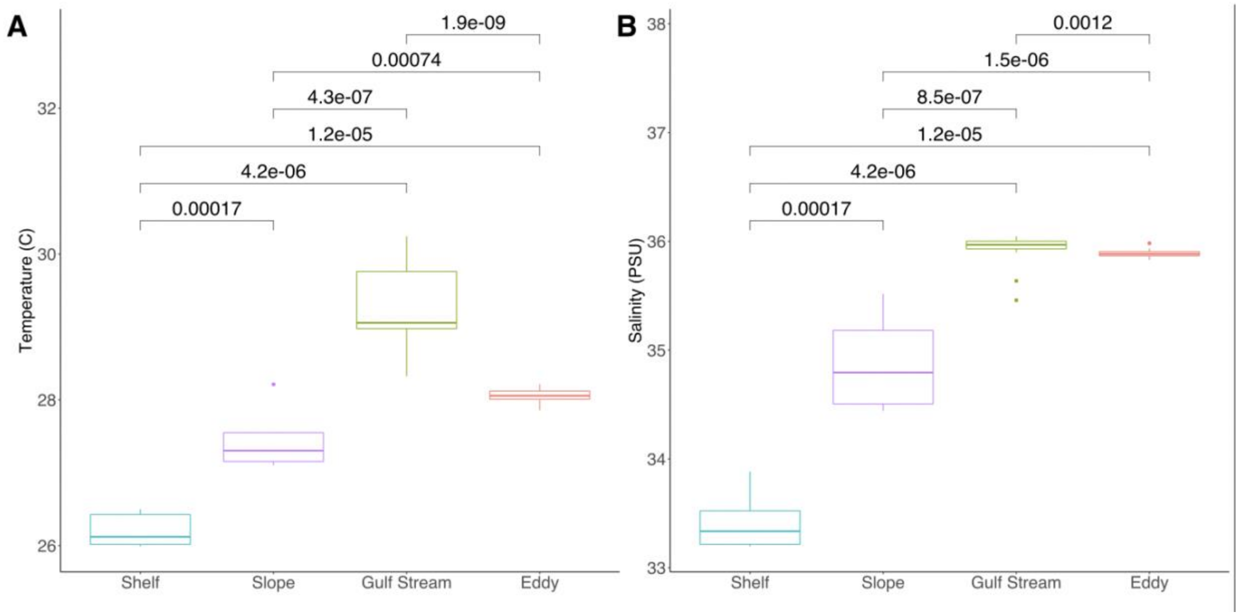


**Figure S1. Environmental variables for physical parameter defined water parcels** (A) Temperature and (B) salinity for each of the temperature and salinity-defined water parcels (Continental Shelf, Continental Slope, Gulf Stream, Eddy) as determined using the flow-through seawater system at ~1m depth. Center lines denotes the median value while the box encompasses the $25^{th}$ to $75^{th}$ percentiles of the dataset. Whiskers denote the $5^{th}$ and $95^{th}$ percentiles and values that fall outside these ranges are shown as individual points. Brackets indicate p-values associated with Wilcoxon Rank Sum test pairwise comparisons.

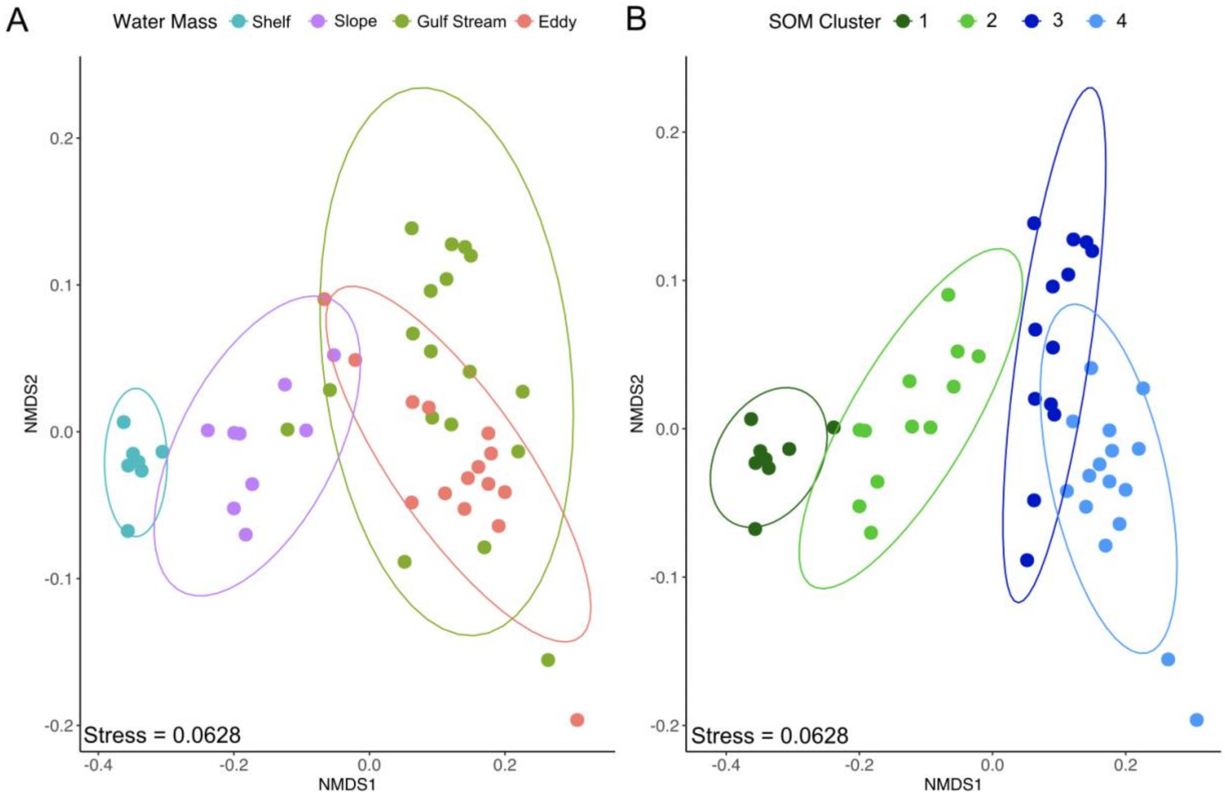


**Figure S2. Non-metric multidimensional scaling (NMDS) ordination comparisons of bacterial communities grouped by physical and biological parameters**. NMDS computed based on Bray-Curtis dissimilarity for 16S rRNA gene libraries and colored by (A) oceanographic water parcels (temperature and salinity) and (B) Self-Organizing Map (SOM) clusters based on the microbiome. Ellipses show the multivariate t-distribution 95% confidence interval for the mean microbial community of each water parcel or SOM cluster.

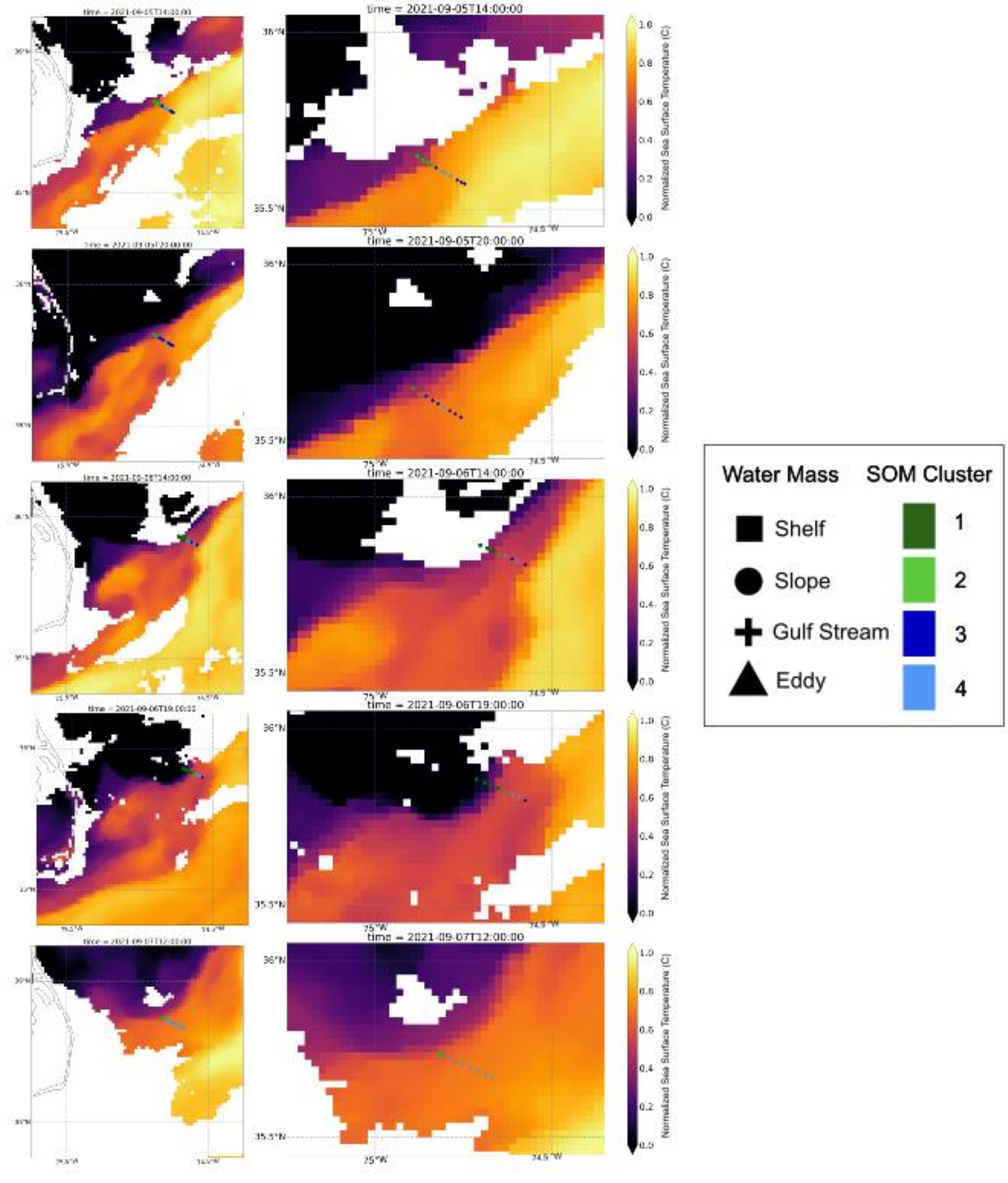


**Figure S3. Maps of sampling sites for five transects off the coast of Cape Hatteras, North Carolina, USA from September 5-7, 2021.** Background color represents sea surface temperatures at time of sampling obtained from the Geostationary Operational Environmental Satellite 16 (GOES-16). Each sampling point is shown along the transects and categorized by oceanographic water parcel (shape), defined by temperature and salinity, and microbial-community based k-means clusters (color), generated using the self-organizing map R package 'kohonen'. Black dotted lines indicate the approximate outline of the frontal eddy during the time of sampling based on a qualitative analysis of the SST imagery.

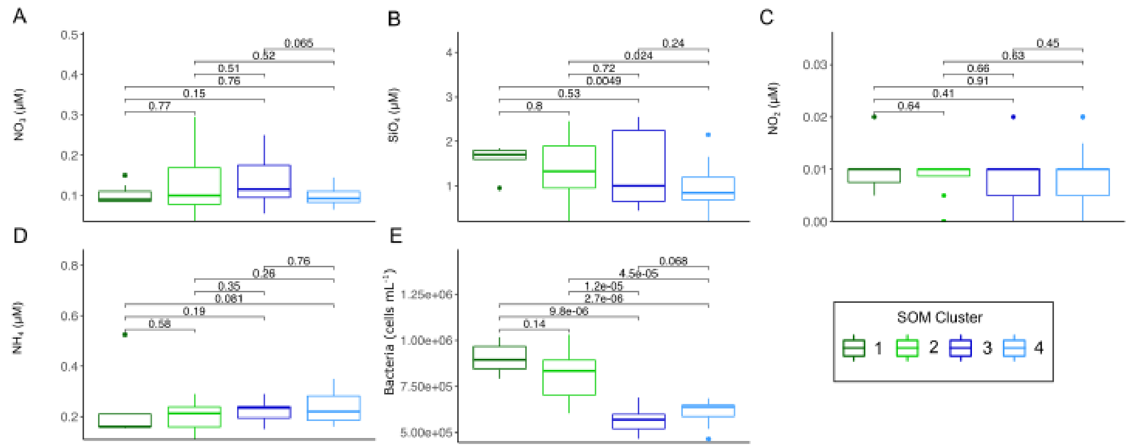


**Figure S4. Additional environmental factors for Self-Organizing Map (SOM) clusters**. Numbered clusters correspond to the most common environments: Continental shelf (Cluster 1), Continental slope (Cluster 2), Gulf Stream (Cluster 3), Eddy (Cluster 4). For the box and whisker plots, the center lines denote the median value while the box encompasses the 25$^{th}$ to 75$^{th}$ percentiles of the dataset. Whiskers denote the 5$^{th}$ and 95$^{th}$ percentiles and values that fall outside these ranges are shown as individual points. Brackets indicate p-values of Wilcoxon Rank Sum pairwise comparisons.

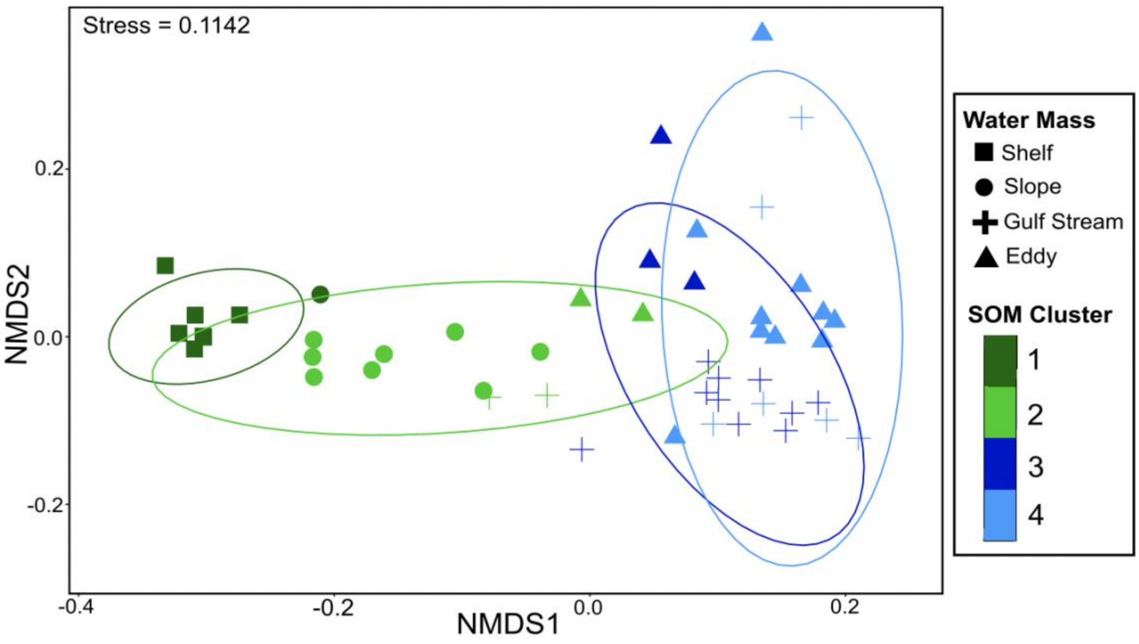


**Figure S5. Non-metric multidimensional scaling (NMDS) ordination computed based on Sørensen similarity for 16S rRNA gene libraries**. Each sampling point is categorized by oceanographic water parcel (shapes), defined by temperature and salinity, and microbial-community based k-means clusters (colors), generated using the self-organizing map R package ‘kohonen’. Ellipses show the multivariate t-distribution 95% confidence interval for the mean of each Self-Organizing Map (SOM cluster).

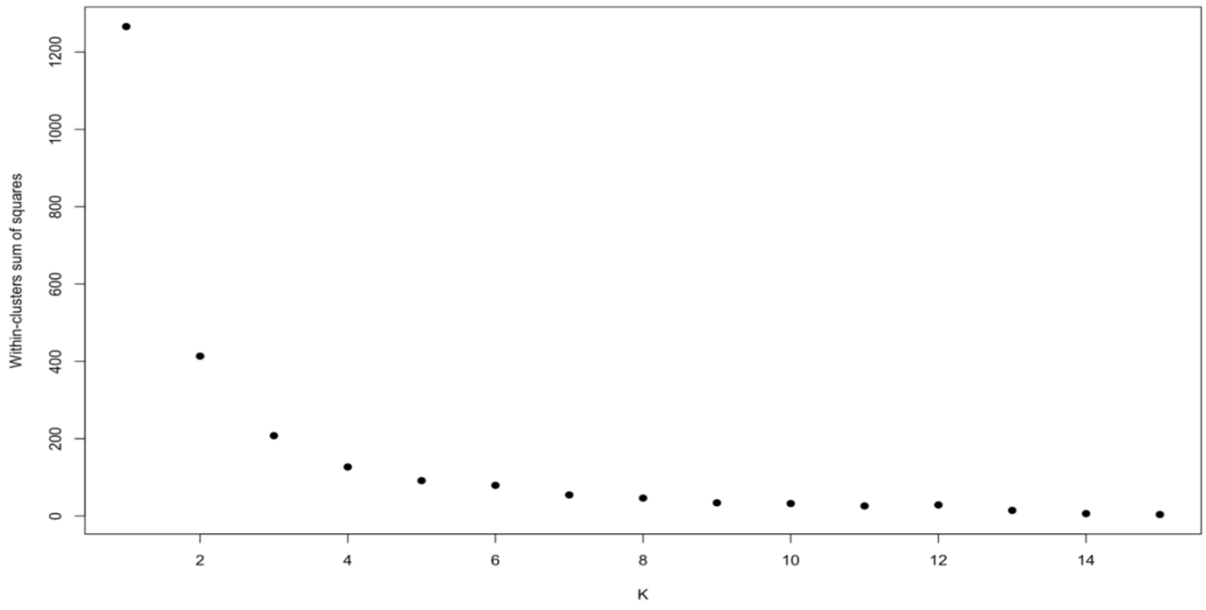


**Figure S6**. **Self-Organizing Map within-clusters sum of squares plot for cluster number selection**. Calculated using a 4x4 grid with hexagonal mapping units in the ‘kohonen’ (v.3.0.11) in R (v4.0.0).
